# Cardiac Specific Overexpression of Transcription Factor EB (TFEB) in Normal Hearts Induces Pathologic Cardiac Hypertrophy and Lethal Cardiomyopathy

**DOI:** 10.1101/2021.02.16.431474

**Authors:** Helena C. Kenny, Eric T. Weatherford, Greg V. Collins, Chantal Allamargot, Taha Gesalla, Kathy Zimmerman, Harsh Goel, Jared M. McLendon, Dao-Fu Dai, Theo Romac, Jennifer Streeter, Jacob Sharafuddin, Abhinav Diwan, Renata Pereira, Long-Sheng Song, E. Dale Abel

## Abstract

TFEB promotes lysosomal biogenesis, autophagy, and lysosomal exocytosis. The present study characterized the consequence of inducible TFEB overexpression in cardiomyocytes in vivo. We generated cardiomyocyte-specific doxycycline inducible (Tet off) mice to achieve spatial and temporal control of TFEB overexpression, by crossing TFEB transgenic mice with mice harboring the tTA transgene (TFEB/tTA). Two weeks after doxycycline removal, an 8-fold increase in TFEB protein expression was observed in transgenic hearts. Heart weight normalized to tibia length was increased by 2.5-fold following TFEB overexpression (TFEB/tTA), characterized by induction of markers of pathological hypertrophy, such as *Nppa, Nppb* and *Acta1*, progressive contractile dysfunction and cardiac dilatation. Overexpression of TFEB resulted in premature death, associated with high degree AV block. Reversal of TFEB overexpression normalized cardiac structure and function. Mitochondrial respiration and ATP levels were preserved after 2-weeks of TFEB induction, despite reduced mitochondrial (OXPHOS) protein expression, mtDNA content, and altered mitochondrial morphology. Signaling through mTOR was induced in TFEB/tTA mice, and when inhibited by rapamycin treatment for 4 weeks, partially offset left ventricular dysfunction. Transcriptome analysis revealed early suppression of mitochondrial metabolic pathways, induction of fibrosis and altered calcium signaling. MCOLN1, a lysosomal calcium release channel, the calcineurin target RCAN1.4, and the mitochondrial calcium uniporter (MCU) were strikingly induced in TFEB/tTA mice. In summary, persistent overexpression of TFEB at high levels (8-fold protein upregulation) in cardiomyocytes promotes pathologic cardiac hypertrophy via suppression of mitochondrial bioenergetic pathways and activation of pro-fibrotic and calcium regulatory pathways.

## Introduction

TFEB, a member of the helix-loop-helix leucine-zipper family of transcription factors, activates the transcription of several autophagic and lysosomal genes by directly binding to the specific 10-base E-box sequence at their promoters [1]. This gene network has been named the Coordinated Lysosomal Expression and Regulation (CLEAR) network [2]. TFEB overexpression increases lysosome number and their degradative ability, and induces the biogenesis of autophagosomes, fusion of autophagosomes with lysosomes and autophagic flux [3]. TFEB directly targets and positively regulates peroxisome proliferator activated receptor-γ coactivator 1α *(Ppargc1α)* and has been shown to regulate energy metabolism in the liver [4] and heart [5, 6]. It also plays a role in mitochondrial autophagy (mitophagy) facilitating parkin-mediated mitochondrial clearance [7]. The regulation of transcription factor EB (TFEB) is mediated through post-transcriptional modifications whereby TFEB localization and activity are mainly controlled by its phosphorylation status. mTOR and ERK2/MAPK1 phosphorylate TFEB at Ser142, retaining it in the cytosol where it remains inactive and bound to the protein chaperone 14-3-3 at the TFEB docking site Ser211 [8]. Conversely, during conditions of stress or starvation, mTOR is released from the lysosomal membrane and becomes inactive [9]. Calcium (Ca^2+^) release from the lysosome through Ca^2+^ channel mucolipin 1 (MCOLN1) activates the phosphatase, calcineurin, which dephosphorylates TFEB and promotes its nuclear translocation [10].

Studies have manipulated the expression of TFEB to assess the effects of autophagy and to investigate the specific role of TFEB in heart failure. Using a model of proteotoxicity (CryAB^R120G^), adenoviral overexpression of TFEB in cultured cardiomyocytes, reduced the accumulation of protein aggregates by increasing autophagic flux [11]. Given that TFEB protein expression was reduced in mouse hearts with proteinopathy, it could represent an important therapeutic target in the setting of cardiac proteotoxicity [12]. However, it remains unclear if TFEB overexpression is beneficial in other physiological and pathological settings. While TFEB is known for its role in lysosomal biogenesis and autophagy regulation, others have reported the importance of TFEB in the regulation of energy metabolism. Transcriptomic analysis in TFEB^-/-^ cardiomyocytes revealed differentially expressed genes involved in nutrient metabolism, cardiac function, DNA repair and damage and apoptosis [5]. Support for the role of TFEB in regulating energy metabolism is also observed in skeletal muscle with TFEB overexpression (AAV-TFEB), which demonstrated that TFEB controls the expression of genes involved in mitochondrial biogenesis, fatty acid oxidation and oxidative phosphorylation [13]. Given the significant role that mitochondria play in calcium homeostasis and considering calcium release from the lysosome is responsible for calcineurin-mediated TFEB dephosphorylation and activation, it is plausible that calcium may also play a key role in TFEB-mediated metabolic regulation. However, the adaptation of the heart to TFEB overexpression in terms of autophagy, mitochondrial morphology and metabolism remains unknown.

TFEB has been advanced as a potential therapeutic target in certain pathological settings. However, additional investigation is required relating to the non-canonical pathways regulated by TFEB, as well as elucidating the specific pathological conditions in which manipulating TFEB could be beneficial. The present study fills an important gap in knowledge regarding the role of sustained activation of TFEB in cardiomyocytes particularly as it relates to autophagy and non-canonical pathways that regulate cardiac mitochondrial metabolism. The present study reveals that 8-fold TFEB overexpression in cardiomyocytes in vivo results in cardiac hypertrophy, left ventricular contractile dysfunction and premature death, likely resulting from high degree AV block. mTOR signaling was induced in TFEB overexpressing hearts and its inhibition by rapamycin treatment partially offset LV dysfunction. Transcriptomic analysis revealed repression of energy metabolism pathways, promotion of pro-fibrotic pathways and altered calcium homeostasis that preceded ventricular dysfunction. Thus, sustained and unregulated TFEB activation stimulates pathways that promote ventricular remodeling.

## Methods

### Animals and animal care

Animal work was performed in accordance with protocols approved by the University of Iowa Animal Care and Use Committee (IACUC). All animals in this study were on the FVB genetic background and equal ratios of both male and female animals were employed. Separate analysis was done to assess the presence of sexual dimorphism. All mice were compared directly to WT controls on the same background and at the same age. TFEB transgenic (Tg) mice were kindly provided by Dr. Diwan, MD, Center for Cardiovascular Research and Division of Cardiology, Washington University School of Medicine. In the Tet-Off expression system, a tetracycline-controlled transactivator protein (tTA), composed of a Tet repressor DNA binding protein (TetR) regulates expression of the target gene, TFEB, which is under control of the transcriptional response promoter element (TRE). The TRE is made up of Tet operator (tetO) sequence fused to a minimal promoter (CMV). In the absence of Doxycycline (DOX), tTA binds to the TRE and activates the transcription of TFEB. TFEB Tg mice were bred with mice expressing tTA under the control of the cardiomyocyte-specific α-myosin heavy chain (αMHC) promoter. TFEB Tg+ or – and αMHC-tTA + or - were used as control littermates which excluded tTA mediated toxicity as a basis for phenotypic disparity. Doxycycline chow (Sigma, St Louis, MO) (1g/kg) was removed from both TFEB/tTA and WT mice at 6-8 weeks old and mice were maintained on a standard chow (2920X Harlan Teklad, Indianapolis, IN, USA). Animals were housed at 22°C with a 12-h light, 12-h dark cycle with free access to water and 2920X chow or doxycycline chow. To assess in vivo mitophagy, TFEB/tTA mice were crossed with mice containing the Mito-Keima transgene, also on the FVB background. The Mito-Keima mice were a kind gift from Dr. Torn Finkle (Center for Molecular Medicine, NHLBI, NIH). Adult cardiomyocytes were isolated from mice with TFEB/tTA plus mito-Keima and were assessed using Leica SP8 confocal microscope. We depict the Mito-Keima fluorescence signal from the excitation at 651 nm (acidic pH) in red and 458 nm (neutral pH) in green. The ratio of the area of lysosomal (red) signal to mitochondrial (green) was used as a measure of mitochondrial delivery to lysosomes. We compared TFEB/tTA/Mito-Keima to WT mice with Mito-Keima as previously described [14].

### Animal Treatments

Rapamycin was obtained from LC Laboratories (Woburn, MA). Rapamycin was dissolved in 2% dimethylsulfoxide (DMSO) 0.2% carboxymethyl cellulose sodium (CMC) and 0.25% tween80 to make a stock solution of 0.2 mg/ml and stored in 1 ml aliquots at −20°C. Vehicle control was made up with the same ingredients except the rapamycin and also aliquoted into 1 ml tubes and stored at −20°C. Each mouse received a dose of 2 mg/kg 5 days/week for 4 weeks by intraperitoneal (i.p) injection. After 4-weeks, mouse tissue was harvested. In a separate cohort of mice, to inhibit autophagosome turnover and to determine autophagic flux, mice were injected with chloroquine diphosphate salt (CQ) (Sigma) after 2-weeks TFEB induction at 48 h (30 mg/kg body weight), 24 h (30 mg/kg), and 2 h (50 mg/kg) prior to harvest.

### Mitochondrial DNA content

Mitochondrial DNA content was quantified by real-time polymerase chain reaction (RT–PCR). Briefly, total DNA was extracted and purified from heart tissue with the DNeasy Kit (Qiagen Inc., Valencia, CA, USA). DNA (5 ng) was used to quantify mitochondrial and nuclear DNA markers. RT–PCR was performed using an ABI Prism 7900HT instrument (Applied Biosystems, Foster City, CA, USA) in 384-well plate format with SYBR Green I chemistry and ROX internal reference (Invitrogen). Analysis of results was automated using scripting with SDS 2.1 (Applied Biosystems), Microsoft Access, and Microsoft Excel. Mitochondrial DNA content *(Cox1)* was expressed relative to genomic *Rpl13a* gene. The following primers were utilized: Primers (5′ to 3′):

- *Cox1*-fwd: gcc cca gat ata gca ttc cc
- *Cox1*-rev: gtt cat cct gtt cct gct cc
- *Rpl13a*-fwd: gag gcc cct acc att tcc ga
- *Rpl13a*-rev: ggc ttc agc cga aca acc tt

### Transmission Electron microscopy

Left ventricle (LV) tissue was isolated and cut into small pieces (<1 mm^3^) and fixed with ½ Strength Karnovsky’s fixative (Final mixture is 2% Paraformaldehyde, 2.5% Glutaraldehyde and 0.1M sodium cacodylate buffer, pH 7.2) overnight at 4 °C and then post-fixed with 1% osmium tetroxide for 1 h. After serial alcohol dehydration (50%, 75%, 95% and 100%), the samples were embedded in Eponate 12 (Ted Pella). Ultramicrotomy was performed and ultrathin sections (70 nm) were post-stained with uranyl acetate and Reynold’s lead citrate. Samples were examined with a JEOL 1230 transmission electron microscope. Mitochondrial volume density, number and size were quantified using Image J.

### WGA staining and stereological quantification

Heart tissue was embedded in paraffin, portioned into 5-μm-thick sections, stained with wheat-germ agglutinin (WGA)–Alexa Fluor 488 conjugate (Invitrogen Corporation, Carlsbad, CA, USA). Images were obtained and processed at the University of Iowa Microscopy Core Facility. From each sample, 10 microscopic fields were analyzed at random, the stage of the microscope being moved blindly. For stereological analysis, cross-sectioned myocytes were selected. Myocyte diameter and number were determined using the Image J.

### Gene expression analysis

Hearts were excised, rinsed in ice-cold phosphate-buffered saline and snap-frozen. Total RNA was extracted from tissues with TRIzol reagent (Invitrogen) and purified using the Direct-zol RNA MiniPrep Plus (Zymo Research). RNA concentration was determined by measuring the absorbance at 260 and 280 nm using a spectrophotometer (NanoDrop 1000, NanoDrop products, Wilmington, DE, USA). Total RNA (50ng/ul) was reverse-transcribed using the High-Capacity cDNA Reverse Transcription Kit (Applied Biosystems, Carlsbad, CA), followed by qPCR reactions using Power SYBR Green (Life Technologies, Carlsbad, CA). Samples were loaded in a 384-well plate in triplicate, and real-time polymerase chain reaction was performed with an ABI Prism 7900HT instrument (Applied Biosystems). The following cycle profile was used: 1 cycle at 95°C for 10 min; 40 cycles of 95°C for 15 s; 59°C for 15 s, 72°C for 30 s, and 78°C for 10 s; 1 cycle of 95°C for 15 s; 1 cycle of 60°C for 15 s; and 1 cycle of 95°C for 15 s. Data were normalized to GAPDH, and results are shown as fold change vs. WT mice. qPCR primers were designed using Primer-Blast [15] and are listed in Supp. Table 1.

### Measurements of Mitochondrial Respiration

Heart fibers were freshly excised and permeabilized with saponin. About 8mg of heart tissue was placed in ice-cold BIOPS (in mM; 7.23 K_2_EGTA, 2.77 CaK_2_EGTA, 20 Imidazole, 0.5 DTT, 20 Taurine, 5.7 ATP, 14.3 PCr, 6.56 MgCl_2_-6H_2_O, 50 MES, pH 7.1 at −20°C). After rapid manual separation of the heart fibers with sharp forceps in BIOPS solution under a microscope, the fibers were quickly placed in a BIOPS and saponin solution (50 ul.ml) and gently agitated at 4°C for 30 minutes. The fibers were then washed in respiration medium three times for 10-minutes (in mM; 1 EGTA, 20 ul blebbistatin, 20 creatine monohydrate, 105 K-MES, 30 KCl, 10 KH_2_PO_4_, 5 MgCl_2_6H_2_0, 5mg/ml BSA, pH 7.4, −20 °C). Fibers were blotted, weighed and immediately placed in the two chambers (OROBOROS, Oxygraph-2k, Innsbruck, Austria) [16] for respiration measurements. Respiration was measured at 30 °C with approximately 3-5mg of permeabilized muscle fibers in each chamber containing 2 ml respiration medium. The software DatLab (Oroboros, Innsbruck, Austria) was used for data acquisition at 2 second time intervals [17]. Respiration was measured using pyruvate/malate (5mM/5mM) or palmitoylcarnitine/malate (0.01mM/5mM), followed by sequential addition of ADP (2.5mM), FCCP (10mM), succinate (10mM), Rotenone (10mM). Oxygen consumption rates were expressed as nmol of O2 × min^−1^ × mg wet fiber weight^−1^.

### Mitochondrial ATP production

The rate of ATP production in the permeabilized fibers was recorded alongside the O2 consumption in real time by monitoring the increase in fluorescence in the respiration chamber coming from NADPH (340ex/460em) using the Horiba Jobin Yvon spectrofluorometer as described [18]. In order to maintain this reaction, 2.5 U/ml glucose-6-phosphate dehydrogenase (Roche), 2.5 U/ml hexokinase (Roche), 5 mM nicotinamide adenine dinucleotide phosphate (NADP+) (Sigma-Aldrich) and 5 mM D-glucose (Sigma-Aldrich) were added to the assay media. P1, P5-Di(adenosine-5’) pentaphosphte (Ap5A) (Sigma-Aldrich) was included in the respiration medium to inhibit adenylate kinase and to ensure that ATP production was solely due to mitochondrial oxidation phosphorylation. An absolute amount of ATP generated across a given time frame was then calculated using a standard curve of fixed concentrations of ATP added to the saturating amounts of hexokinase, glucose, G6PDH, and NADP^+^.

### Histology

Mouse hearts were fixed in 4% paraformaldehyde (PFA), embedded in paraffin, proportioned into 5-μm sections, and stained with hematoxylin and eosin, Masson’s trichrome and Picrosirus Red (PSR) under standard protocols. Images were obtained using the Olympus BX-61 microscope and Leica Aperio Ariol Digital Slide Scanner (Leica Biosystems). Picrosirius red–stained sections were examined using polarizing microscopy and a color wavelength (from yellow-orange to deep red) indicative of collagen fiber thickness was noted in collagen containing regions. For immunofluorescence, heart sections were incubated with rabbit anti-TFEB (SC48784, 1:500 dilution) antibody for 1 h and subsequently Alexa Fluor 568 goat anti-rabbit (Invitrogen, A-21069) or Alexa Fluor 488 goat anti-mouse (Invitrogen, A11017) secondary antibody (1:500 dilution) for 1 h at room temperature. TFEB was detected with excitation/emission of 543/618 nm by indirect fluorescence on Leica SP836 Confocal microscope.

### Echocardiography

Transthoracic echocardiograms were acquired from sedated (midazolam 0.1 mg subcutaneous injection) mice. The anterior chest hair was removed with Nair (Church & Dwight) and warmed gel was applied. The mouse was held gently by the nape of the neck and positioned in the left lateral position. Cardiac images were obtained via a 30-MHz linear array transducer applied to the chest. The transducer was coupled to a Vevo 2100 imager (Visual Sonics). Images of the short and long axis were obtained with a frame rate of ~180–250Hz. All image analysis was performed offline using Vevo 2100 analysis software (v.1.5). Endocardial and epicardial borders were traced on the short axis view in diastole and systole. LV length was measured from endocardial and epicardial borders to the LV outflow tract in diastole and systole. The biplane area–length method was then performed to calculate LV mass and ejection fraction.

### Electrocardiography

Mice were anesthetized with 2% isoflurane and placed on a surgical monitoring board to collect EKG and maintain body heat (Indus, Rodent Surgical Monitor+). Three-Lead EKG was acquired using external titanium leads placed subcutaneously on limbs. Signals were routed through an analog output device (Indus, #351-0066-01) and converted into digital signals for analysis (AD Instruments, Powerlab 8/8, Labchart Pro v8.1.13, EKG module v2.4). Time selection windows of 10 seconds were used for EKG analyses with 4 beat averaging with standard mouse detection settings, and without excluding “outlier” beats. Data were exported for offline analyses using Microsoft Excel and Graphpad Prism v8.2.1.

### Western blotting

Approximately 50mg of heart tissue were lysed in 500ul RIPA buffer containing (in mmol/l) 50 HEPES, 150 NaCl, 10% glycerol, 1% Triton X-100, 1.5 MgCl_2_, 1 EGTA, 10 sodium pyrophosphate, 100 sodium fluoride, and 100 μmol/l sodium vanadate. Right before use, HALT protease/phosphatase inhibitors (Thermo Fisher Scientific) were added to the lysis buffer and samples were processed using the TissueLyser II (Qiagen Inc.). Tissue lysates were resolved on 12% SDS–PAGE, electrophoresed at 60V for 16 hours, then transferred to PVDF membranes (Millipore Corp). Membranes were incubated with primary antibodies overnight and with secondary antibodies for 1 h. Western blots were performed using different antibodies at 4°C overnight (Supp. Table 2)

### RNA sequencing

To determine the changes in functional and metabolic pathways, transcriptome analysis was performed in cardiac specific TFEB/tTA and WT) mice after 1-week doxycycline removal. 1-week timepoint was chosen as the mice still had preserved LV function. Transcriptome analysis was performed in ventricular samples from 5 female hearts per group. Sequencing libraries were prepped using the Illumina TruSeq Stranded Total RNA kit and sequenced on a NovaSeq 6000. Reads were aligned, mapped, and quantified using HISAT2 (v2.2.1) and featureCounts (version 2.0.1) with differential expression analysis using DESeq2 (v1.30.0) [19–21]. Alternatively, reads were pseudo aligned and quantified using Kallisto (v0.46.2) followed by differential expression analysis with DESeq2 [22]. Set-based enrichment analysis was performed using the R (version 4.0.2) statistical software environment Enrichment Browser package (v2.20.4) and the Pathway Analysis with Down-weighting of Overlapping Genes (PADOG) method [23, 24]. Additional pathway analysis was performed with Ingenuity^®^ Pathway Analysis (IPA^®^) software from Qiagen. Heatmaps and plots were produced using pheatmap and ggplot2 packages in R.

### Data analysis

All data are reported as mean ± SEM. Student’s *t*-test was performed for comparison of two groups, and ANOVA followed by Tukey multiple comparison test was utilized when more than three groups were compared. A probability value of *p* ≤ 0.05 was considered significantly different. Statistical calculations were performed using the GraphPad Prism software (La Jolla, CA, USA)

## Results

### Pathological cardiac hypertrophy develops following TFEB induction in cardiomyocytes

TFEB was overexpressed in cardiomyocytes using a doxycycline inducible Tet-off model (Fig. 1A). At 6-8 weeks of age, DOX was removed and an 8-fold increase in TFEB protein expression (Fig. 1B) that paralleled induction of *Tfeb* mRNA was observed after two weeks (Supp. Fig 1A). Western blot analysis of skeletal muscle and lung tissues showed no change in TFEB protein expression, confirming selective TFEB overexpression in the heart (Supp. Fig 1B-C). Immunofluorescence imaging showed increased TFEB colocalization in the nuclei of cardiomyocytes (Supp Fig. 1D-E), which appeared enlarged relative to nuclei negative for TFEB (Supp. Fig 1F). Terminally differentiated cardiomyocytes will increase in size, not number, in response to physiological or pathological stress [25]. TFEB overexpression increased heart weight normalized to tibia length (HW/TL) after 1- and 2- and 25-weeks after TFEB induction (Fig. 1C). This coincided with a reduction in lung weight normalized to tibia length at weeks 2 and 25-weeks post-induction (Fig. 1D). Atria weight/tibia length was also increased after 25-weeks (Supp. Fig. 1G and Fig. 1E). Wheat Germ Agglutinin (WGA) staining revealed a significant increase in cardiomyocyte size (Fig. 1F-G). qPCR analysis revealed increased mRNA expression of markers of pathological hypertrophy after 1- and 2-weeks post-induction (Supp Fig 1H and Fig. 1H).

**Figure 1.**
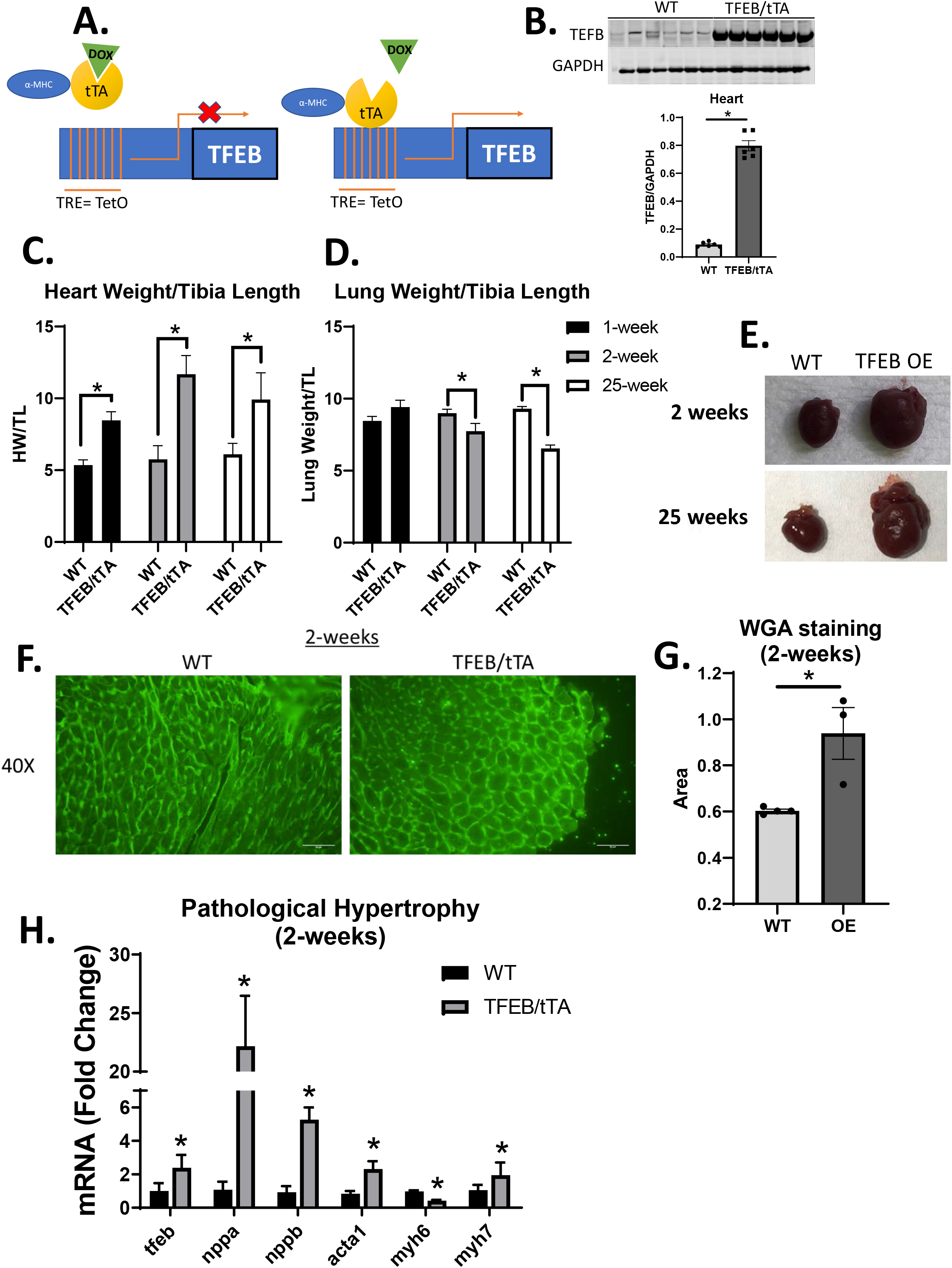
Pathological cardiac hypertrophy develops following TFEB induction in cardiomyocytes. Tet-off model schematic illustrating generation of the cardiac specific TFEB Tg mice crossed with alpha-MHC tTA with or without doxycycline (1A), representative western blot for TFEB (Bethyl, 65KDa) in WT and TFEB Tg heart lysates 2 weeks after doxycycline removal (n=6) (1B), heart weight/tibia length (HW/TL), lung weight/tibia length of WT and TFEB/tTA hearts after 1, 2- and 25-weeks doxycycline removal (1C). Images of WT heart and TFEB/tTA heart after 2- and 25-weeks doxycycline removal (1D), cross sections of heart tissue from mice two weeks after doxycycline removal stained with wheat-germ agglutinin (WGA). Scale bar = 20 μm (n=3-4) (1E), cardiomyocyte area quantified from WGA staining using Image J (1F), mRNA relative expression of pathological markers of hypertrophy (nppa, nppb, acta1, myh6, myh7) in WT and TFEB/tTA mouse hearts (n=6/group) (1G), *p<0.05 WT versus TFEB/tTA using Student’s T-test, graphs represent mean ± SEM.

### Heart failure and premature mortality following TFEB overexpression in cardiomyocytes

Cardiac overexpression of TFEB induced left ventricular dysfunction. After 1-week of TFEB induction, cardiac function was preserved (Supp. Fig 2A); however, 2-weeks after DOX removal, reduced ejection fraction (EF%), a significant increase in end-diastolic volume and end-systolic volume, mass (g) and volume/mass were evident (Fig. 2 A-F). LV dysfunction progressed over 23 weeks confirming the presence of heart failure with reduced ejection fraction (HFrEF). A separate analysis showed no significant difference in LV dysfunction between male and female mice indicating no sexual dimorphism (Supp. Fig. 2B-C). Cardiac specific over expression of TFEB also increased mortality with a 50% death rate by 25 weeks following doxycycline removal (Fig. 2G). A striking induction of transcriptional activation of cardiac fibrosis after 1-week (Supp. Fig. 3A), did not correlate with increased cardiac fibrosis by week-2 as evidenced by picrosirius red and Trichrome staining (Supp. Fig. 3B-C). The dynamic fibrosis/remodeling response is supported by qPCR analysis of fibrosis markers at 2- weeks showing normalization of these pro-fibrotic transcripts (Supp. Fig. 3E).

**Figure 2.**
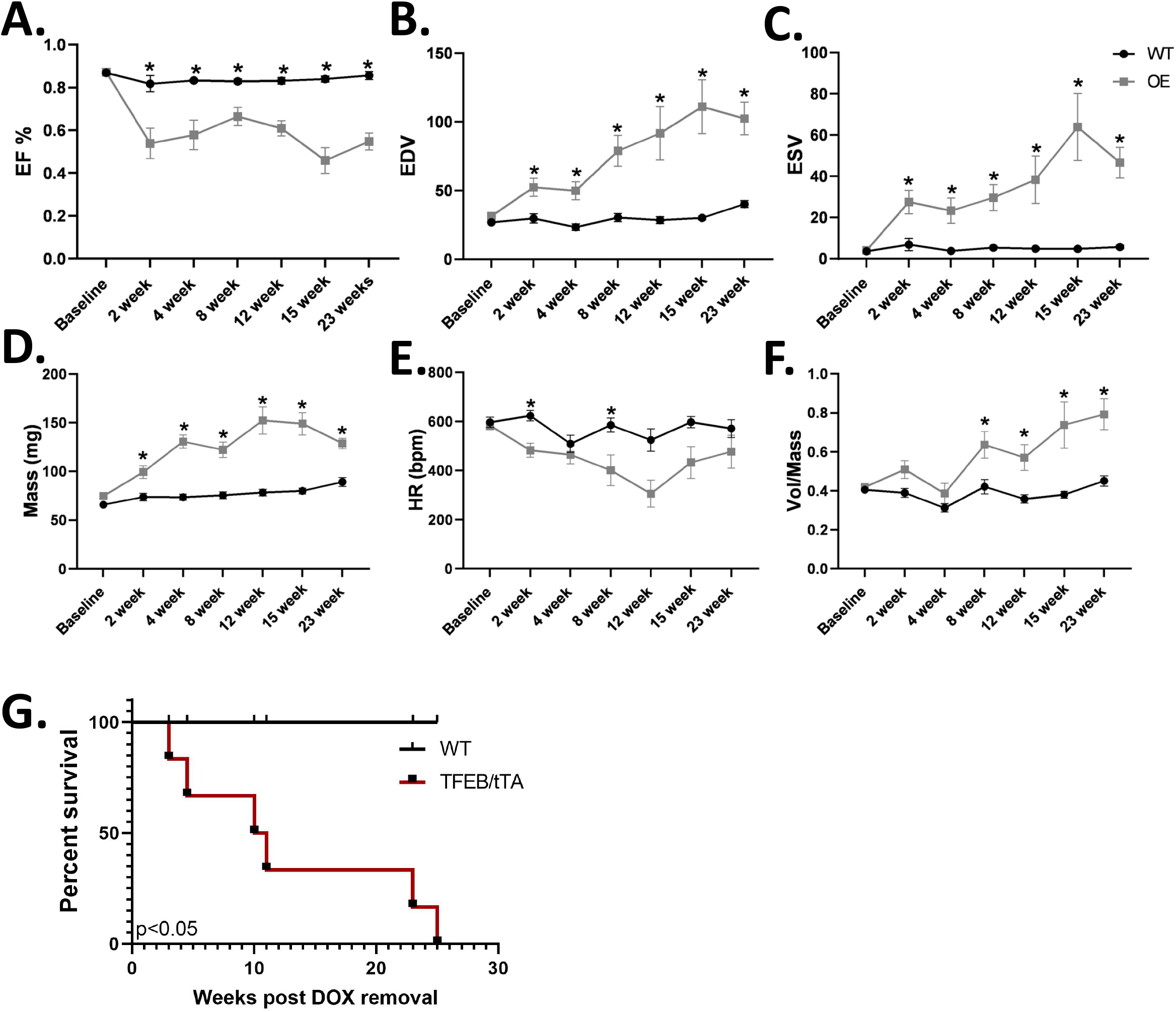
Heart failure and premature mortality following TFEB overexpression in cardiomyocytes. Echocardiography data depicting ejection fraction (EF%), end-diastolic volume (EDV), end-systolic volume (ESV), mass, heart rate (HR) in beats per minute (bpm), volume normalized to mass (volume/mass), at baseline, 2, 4, 8, 12, 15- and 23-weeks post doxycycline removal in WT and TFEB/tTA mice, (WT n= 9-16 and TFEB/tTA n= 6-11) (2A-F), survival curve for WT and TFEB/tTA mice showing percent survival over time (weeks) (2G). *p<0.05 WT versus TFEB/tTA using Student’s T-test, graphs represent mean ± SEM.

### Reversibility of heart failure following reversal of TFEB overexpression

In two separate cohorts, doxycycline was removed for 3 or 5 weeks to induce TFEB overexpression, which led to LV dysfunction. These mice were then put back on doxycycline to suppress TFEB protein expression for an additional 3 or 5 weeks, respectively. The observed cardiac dysfunction after three and five weeks was reversed by reintroducing doxycycline for either 3 or 5 weeks (Fig. 3A-D and Supp. Fig 4.A-D). Cardiac hypertrophy was largely attenuated upon DOX re-introduction, although an increase in HW/TL persisted (Supp. Fig. 4F and Fig. 3E). However, the induction of transcriptional markers of pathological LVH was normalized (Fig. 3H). Thus, the adverse consequences of cardiomyocyte TFEB overexpression can be reversed by normalizing TFEB protein levels.

**Figure 3.**
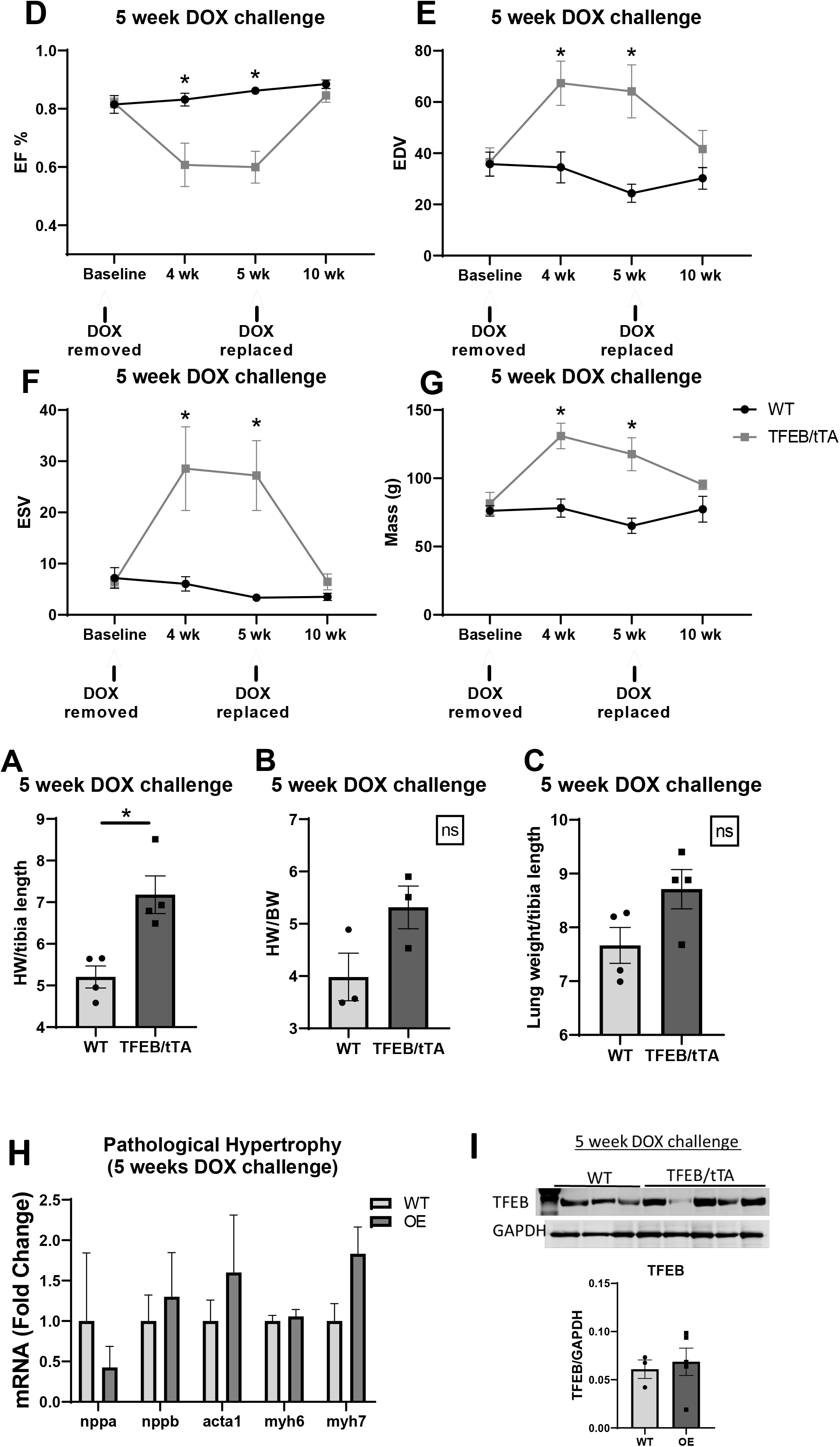
Reversibility of heart failure following reversal of TFEB overexpression. Heart weight normalized to tibia length (HW/TL), heart weight normalized to body weight and lung weight normalized to body weight in WT and TFEB/tTA hearts after removal of doxycycline for 5-weeks and replacement of doxycycline for a further 5-weeks, (n=3-4/group) (3A-C). Echocardiography on WT and TFEB/tTA mice at baseline, 4-weeks after doxycycline removal, at 5-weeks and 10-weeks, ejection fraction (EF%), end-diastolic volume (EDV), end-systolic volume (ESV), mass (g) (WT n=5 and TFEB/tTA n=6) (3D-G), mRNA relative expression of pathological markers of hypertrophy (nppa, nppb, acta1, myh6, myh7) in WT and TFEB/tTA mouse hearts after 10 weeks (5-week off DOX and 5-weeks back on DOX) (3H), Western blot for TFEB (Bethyl, 65KDa) in WT and TFEB Tg heart lysates after 10-weeks (WT n=3, TFEB/tTA n=5) and quantification by densitometry (3I). *p<0.05 WT versus TFEB/tTA using Student’s T-test and two wat ANOVA, graphs represent mean ± SEM.

### TFEB overexpression in cardiomyocytes inhibits autophagy and mitophagy

Autophagic flux was determined by LC3 and p62 immunoblotting after injection of chloroquine (CQ). CQ neutralizes lysosomal pH and impairs the fusion between autophagosomes and lysosomes. Increased LC3-II accumulation after CQ injections is indicative of increased autophagic flux [26]. CQ increased LC3 I accumulation, with no change in LC3 II and a reduction in LC3II/I ratio in TFEB/tTA hearts compared to WT treated with saline. A significant increase in the accumulation of p62 in TFEB/tTA hearts compared to WT treated with saline was also observed (Fig. 4D-H). The significant reduction in LC3II/I ratio together with an increase in p62 in TFEB/tTA in vehicle-treated mice suggests reduced autophagic flux in TFEB overexpressing hearts. Furthermore, we observed no change in LC3II or LC3 II/I ratio in CQ-treated TFEB/tTA mice, further supporting the conclusion that autophagic flux is reduced in TFEB/tTA hearts (Fig. 4D-H). ULK1 phosphorylation of the mTOR target Ser757 was increased, that could also lead to autophagy inhibition (Supp. Fig. 5A-C).

**Figure 4.**
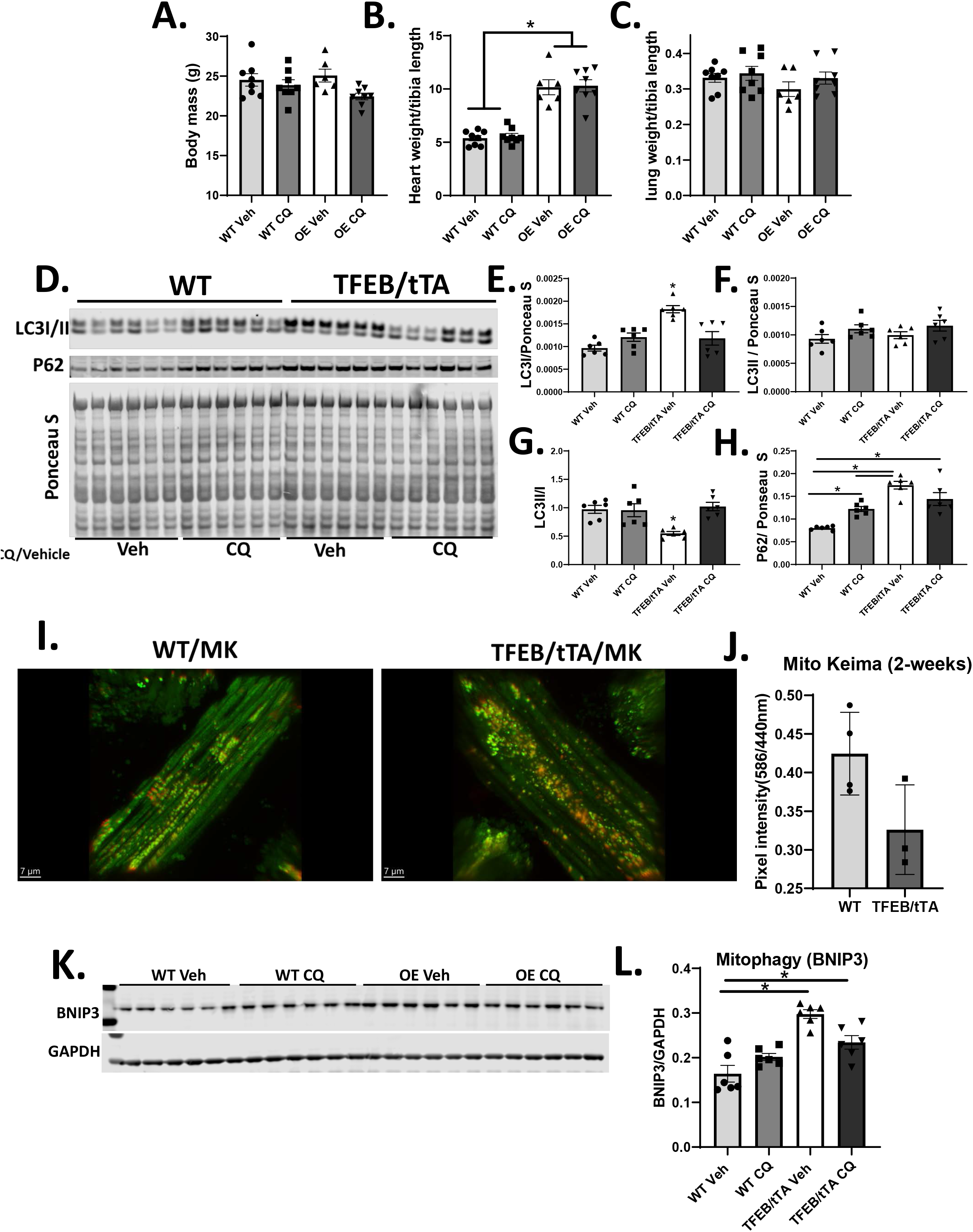
TFEB overexpression in cardiomyocytes inhibits autophagy and mitophagy. Body mass (g), heart weight/tibia length (HW/TL) and lung weight/tibia length in WT and TFEB/tTA (OE) treated with Vehicle and chloroquine (CQ) over 24 hours (4A-C), western blot images TFEB, LC3B, P62 and ponceau stain in WT Veh, WT CQ, OE Veh and OE CQ heart lysates after 24-hour treatment, quantification by densitometry of LC3I, LC3II normalized to ponceau stain, ratio of LCII/I, P62/ponceau S (4D-H). Merged image of WT and TFEB/tTA mice expressing the fluorescent protein keima; keima localized to the inner mitochondrial membrane has an excitation spectrum that changes according to pH (shorter wavelength (458nm) at neutral environment (mitochondria, pH 8) and longer wavelength (561nm) is an acidic environment (lysosome, pH 4.5). Quantification expressed as a ratio of pixel intensity (561nm/458nm) (n=3/group) (4I-J), western blot images for BNIP3 in in WT Veh, WT CQ, OE Veh and OE CQ heart lysates after 24-hour CQ and vehicle treatment, quantification by densitometry and normalized to GAPDH (4K-L). Student’s T-test and two-way ANOVA were used for statistical analysis, graphs represent mean ± SEM.

To assess the effect of TFEB overexpression on mitophagy, TFEB/tTA were crossed with mice that express Mito-Keima. Mito-Keima exhibited a clear pH dependent fluorescence. The ratio of 561nm/458nm (red/green) fluorescence in WT mouse hearts was increased relative to TFEB/tTA hearts (Fig. 4I-J and Supp. Fig 5D). Although BNIP3 levels were increased in vehicle treated TFEB overexpression hearts relative to vehicle-treated WT, no further increase in BNIP3 was observed in CQ treated TFEB/tTA hearts (Fig. 4K-L). Together, these data suggest that an 8-fold increase in TFEB protein expression after 2-weeks of TFEB induction inhibits autophagy and mitophagy.

### Inhibition of mTOR signaling partially attenuates heart failure and hypertrophy induced by TFEB overexpression

Cardiomyocyte growth is generally accompanied by increased protein synthesis. mTORC1 activation promotes ribosomal protein production (mRNA translation) by activating ribosomal protein S6 kinase and suppresses autophagy in part by inhibiting ULK1. mTOR also phosphorylates TFEB, rendering it inactive and retaining it in the cytosol. mTOR signaling was induced 1- and 2 weeks after TFEB induction in TFEB/tTA hearts relative to WT (Supp. Fig. 6A and Fig. 5A-E). Rapamycin treatment for 4-weeks after removal of doxycycline reduced p-mTOR and p-S6 protein (Fig. 5I) but did not reduce cardiac hypertrophy in TFEB/tTA mice (Fig. 5G). Nonetheless, the reduction in lung weight/tibia length evident in the TFEB/tTA group treated with vehicle, was normalized by rapamycin treatment (Fig. 5H). Rapamycin partially attenuated left ventricular dysfunction in TFEB/tTA mice compared to WT (Fig. 5 J-M). These data suggest that mTOR activation could contribute to pathological remodeling in TFEB overexpressing mice and might also contribute to autophagy inhibition.

**Figure 5.**
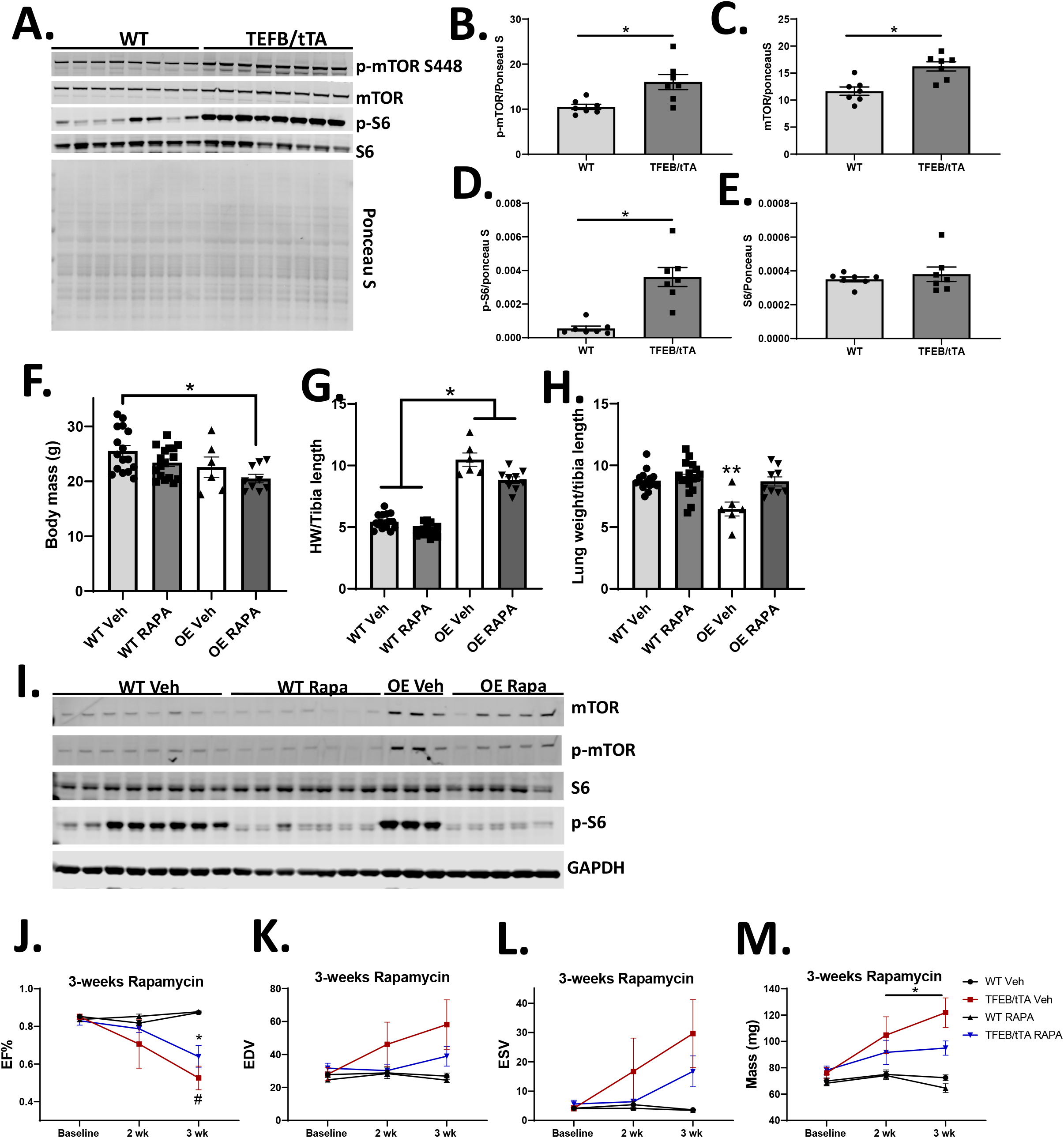
Inhibition of mTOR signaling partially attenuates heart failure and hypertrophy induced by TFEB overexpression. Western blot images (p-mTOR, t-mTOR, p-S6, S6 and ponceau stain) in WT and TFEB/tTA heart lysates after 2-weeks (WT n=8, TFEB/tTA n=8) (5A), quantification of western blots by densitometry and normalized to total protein (ponceau S) for p-mTOR, t-mTOR, p-S6, S6 (5B-E). Body mass, heart weight/tibia length (HW/TL) and lung weight/tibia length in WT and TFEB/tTA (OE) mice treated for 4-weeks with vehicle or rapamycin (WT Veh, WT Rapa, OE Veh and OE Rapa) (5F-H). Western blot images (p-mTOR, t-mTOR, p-S6, S6 and gapdh) in WT Veh, WT Rapa, OE Veh and OE Rapa heart lysates after 4 weeks of vehicle or rapamycin treatment (5I). Echocardiography in WT Veh, WT Rapa, OE Veh and OE Rapa mice at baseline and after 2- and 3-weeks rapamycin treatment, ejection fraction (EF%), end-diastolic volume (EDV), end-systolic volume (ESV), mass (g) (5J-M). *p<0.05 WT versus TFEB/tTA using Student’s T-test and repeated measures ANOVA for echo data, graphs represent mean ± SEM.

### RNA-seq analysis of mRNA isolated from mice following overexpression of TFEB for one week

Because TFEB may have broad transcriptional targets, we conducted transcriptomic analysis to identify mechanisms that if present at the onset of ventricular dysfunction (Supp. Fig. 2A), could inform the pathophysiology of heart failure in this model of TFEB overexpression. Genes that were differentially regulated were filtered based on 1.5-fold changed and an adjusted p value of 0.05. Overall, relative to WT mice, TFEB overexpression resulted in an increase in genes that are positive regulators of cardiac hypertrophy, together with a reduction in genes that mediate cardiac contraction (Fig. 6A). Calcium ion regulatory genes were altered, suggesting altered calcium homeostasis (Fig. 6A). A striking increase in fibrosis genes after one-week was evident (Fig. 6A). There was a broad repression of mitochondrial energy metabolism as evidenced by reduced expression of mitochondrial electron transport chain complex genes (Fig. 6A), tricarboxylic acid cycle, pyruvate metabolism and fatty acid β oxidation (Supp. Fig 7A-C). The most highly regulated canonical pathways, their upstream regulators and disease pathways that were altered in response to TFEB overexpression are shown in Fig. 6B-C. This analysis supports induction of pro-fibrotic regulators and repression of upstream modulators of mitochondrial metabolism.

**Figure 6.**
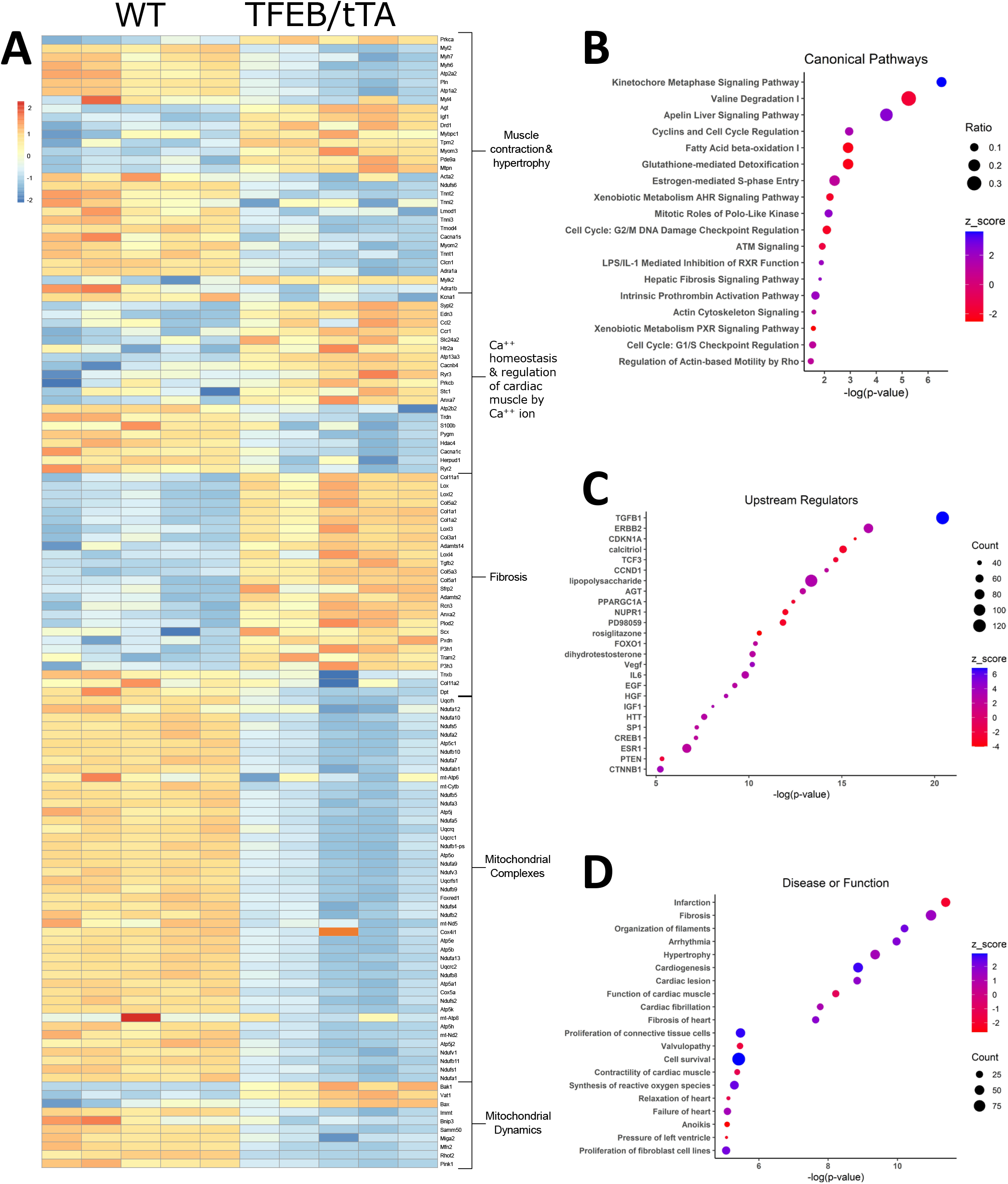
RNA-seq analysis of mRNA isolated from mice following overexpression of TFEB for one week. Heatmap of log2(x+1) normalized counts of differentially expressed genes grouped into pathways or gene sets identified by enrichment analysis performed using Enrichment Browser and the Pathway Analysis with Down-weighting of Overlapping Genes (PADOG) algorithm in WT and TFEB/tTA hearts (n=5/group) (6A). Ingenuity pathway analysis of differentially expressed genes for canonical pathways, upstream regulators, and disease or function (6B-D). Canonical pathways enriched in the differential expression analysis plotting the log(p-value), relative number of genes differentially expressed in the data set relative to the number of genes in the pathway (ratio) and z-score which indicates the predicted activation (positive) or inhibition (negative) of the pathway (6B). Upstream regulators enriched in the data set. Count indicates the number of genes differentially expressed in the data set annotated as regulated by the given factor (6C). Disease or function annotations enriched in the differential expression analysis (6D). Data are filtered based on fold change (1.5-fold) and for an adjusted p value of 0.05.

### Mitochondrial morphology and protein content following 2 weeks of TFEB overexpression in the heart

Growing evidence suggests that TFEB plays a role in mitochondrial quality control [27]. Given our RNASeq data suggesting changes in mitochondrial pathways, we assessed changes in mitochondrial morphology and function. Analysis of TEM micrographs in TFEB/tTA hearts reveal an increase in mitochondrial number together with a reduction in cristae index score, indicating reduced cristae integrity (Fig. 7A-C). There were no changes in other mitochondrial morphological parameters investigated, such as circularity index, aspect ratio, length or width (Sup. Fig. 7A-E). Peroxisome proliferator-activated receptor gamma coactivator 1-alpha (PGC1α) protein and mRNA levels were induced in TFEB/tTA mice, which could contribute to increased mitochondrial number after 2-weeks of TFEB induction (Fig 7. D-E and Supp. Fig. 8A-E). Despite increased mitochondrial number, there was a reduction in mtDNA at 2-weeks (Fig 7F). Altered mitochondrial dynamics was evident with increased protein expression of mitofusin 1 and 2 (MFN1/2), YME1L and total GTPase dynamin-related protein 1 (DRP1) (Fig7G-M). Proteins involved in oxidative phosphorylation were repressed in TFEB/tTA hearts relative to WT (Fig 7N-O), nonetheless, pyruvate-supported mitochondrial respiration, ATP production and P:O ratio were preserved after 2-weeks of TFEB induction (Supp. Fig. 8G-I). The endoplasmic reticulum (ER) has a close relationship with the mitochondria and forms contact sites with mitochondria (MERCs), a tethering point separated by a narrow cleft (10-80nm) which allows for the exchange of lipids and calcium [28]. Importantly, an increase in markers of ER stress signaling (p-eif2α, PERK and IRE1α) was observed in TFEB/tTA hearts relative to WT after 2-weeks (Supp. Fig 9A-F).

**Figure 7.**
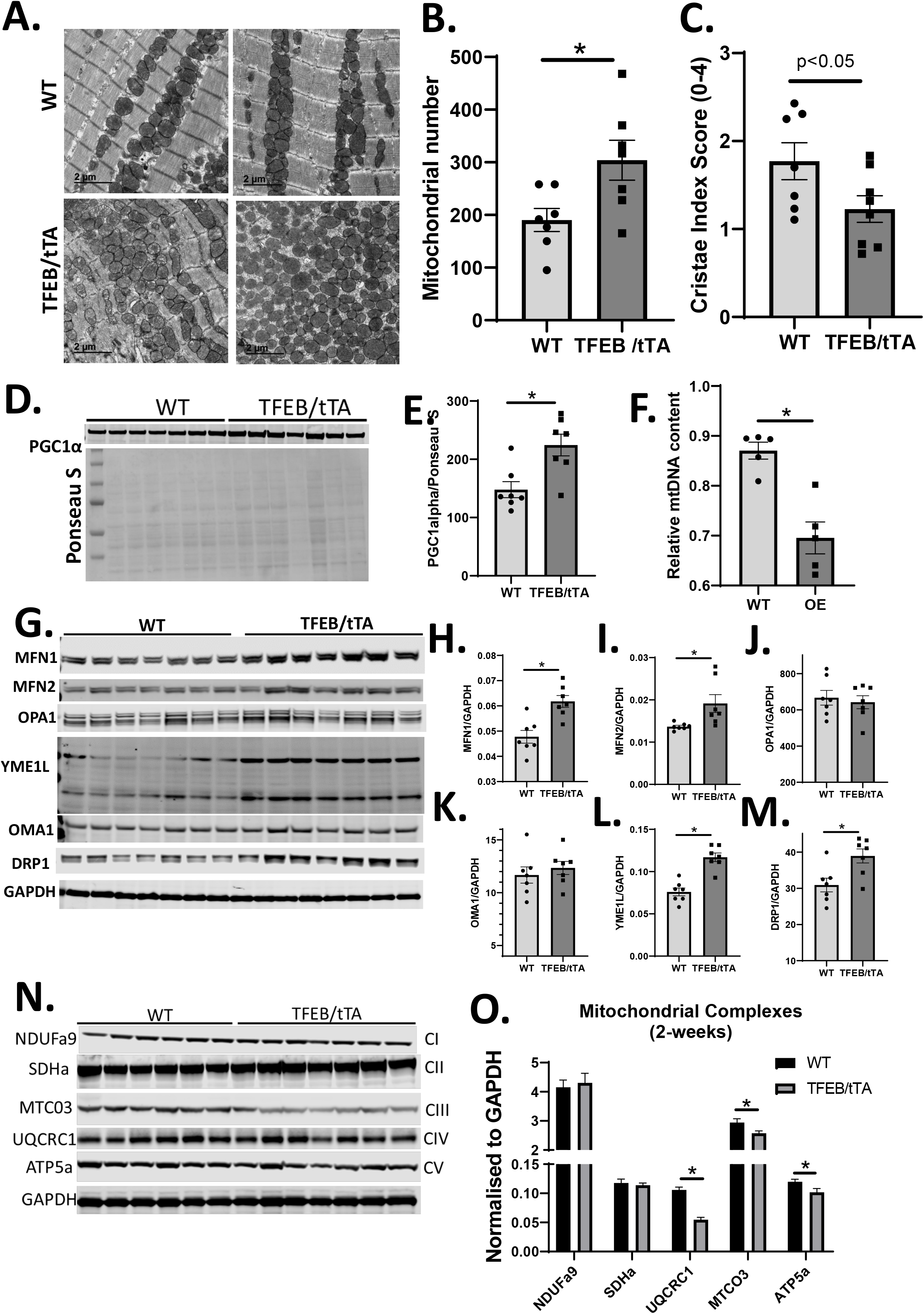
Mitochondrial morphology and protein content following 2 weeks of TFEB overexpression in the heart. Transmission electron microscopy (TEM) in WT and TFEB/tTA hearts (2.5K magnification and 2uM scale bar), quantification of TEM parameters using image J; mitochondrial number and cristae index score (1-4) (7A-C). Western blot image of Peroxisome proliferator-activated receptor-gamma coactivator (PGC)-1alpha and ponceau stain in WT and TFEB/tTA heart lysates (n=7/group), quantification of PGC1-alpha western blot normalized to ponceau (7D-E).Relative mitochondrial DNA content in WT and TFEB/tTA hearts (7F). Western blot images for mitochondrial dynamics proteins (Mfn1, Mfn2, OPA1, Yme1L, OMA1, DRP1 and GAPDH (n=7/group), quantification of western blots using densitometry, Mfn1, Mfn2, Opa1, Yme1L, Oma1, Drp1 (7-G-M). Western blot images of mitochondrial complex proteins (NDUFa9, SDHa, MTCO3, UQCRC1, ATP5a and GAPDH) in WT and TFEB/tTA heart lysates (n=6-7), western blot quantification by densitometry of mitochondrial complex proteins (7N-O). Student’s T-test and two-way ANOVA were used for statistical analysis, graphs represent mean ± SEM.

### Altered calcium regulatory proteins and arrhythmias in TFEB overexpressing hearts

TFEB is regulated by the phosphatase, calcineurin which is activated by lysosomal calcium. Expression levels of mucolipin 1 (mcoln1), a lysosomal calcium releasing channel was increased by TFEB overexpression (Fig. 8A). Increased lysosomal calcium leak is coupled to increased expression of mcip 1.4, a known calcineurin interacting protein (Fig. 8B). Additionally, mitochondrial calcium uniporter (MCU) protein expression is upregulated (Fig. 8C). Together, these changes point to altered calcium regulation in TFEB/tTA hearts. TFEB/tTA mice exhibit premature atrial contractions, atrial flutter and first-degree atrioventricular block after 4-weeks and a high degree AV block at 25 weeks which may contribute to their premature mortality (Fig. 8F-G and Supp. Fig 10 A-D). Transcriptional analysis further supports the hypothesis that regulation of calcium homeostasis is altered in TFEB/tTA mouse hearts compared to WT mice. The cacna1c gene encodes subunits of the calcium channel Cav1.2. Cacna1c gene expression repressed in this current model (Fig. 6A) and impairment in cellular calcium uptake is implicated by the reduction in Cav1.2 protein expression in TFEB/tTA compared to WT after 2-weeks (Fig. 8D-E). We also observed SR calcium gene changes with reduction in ryr2 and trdn gene expression.

**Figure 8.**
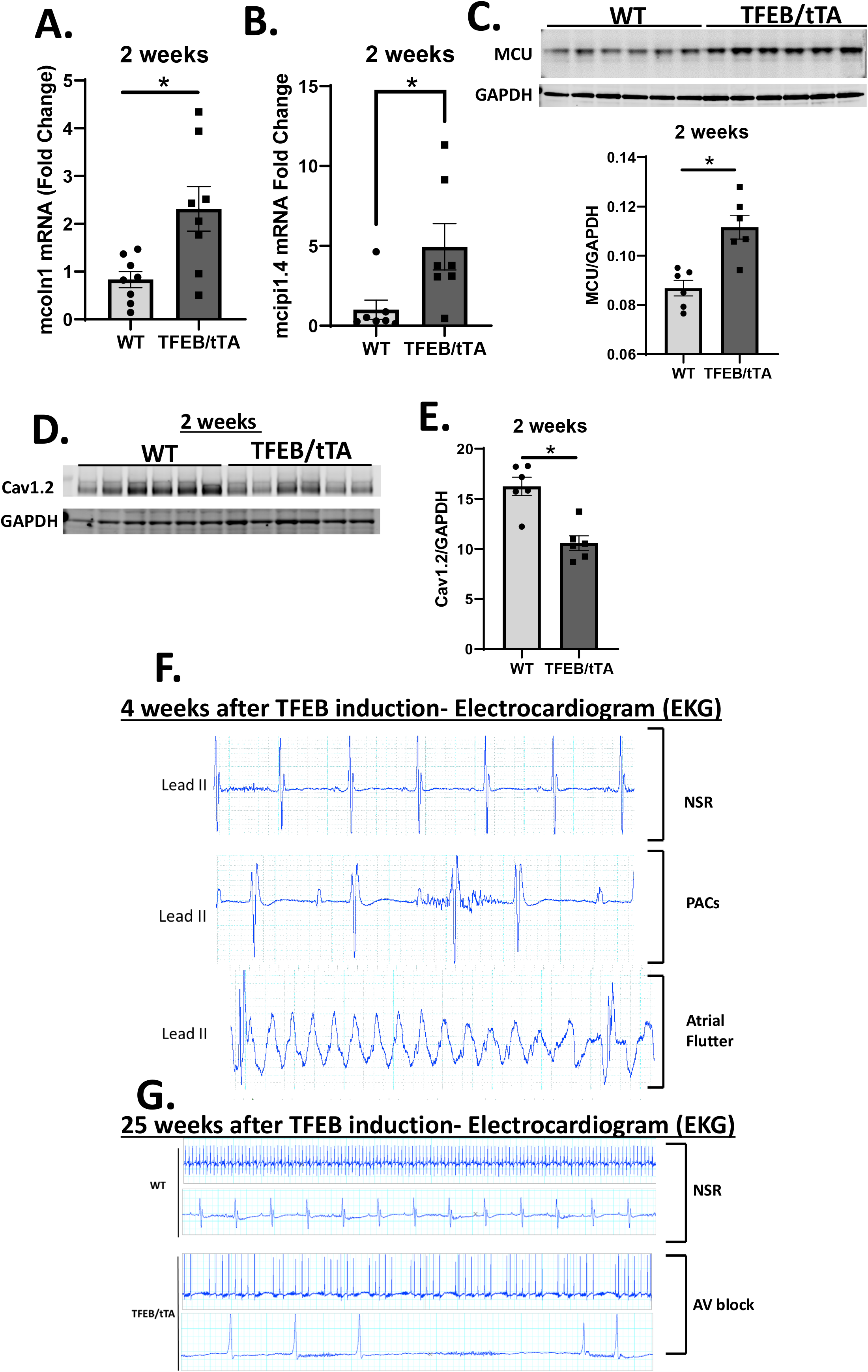
Altered calcium regulatory proteins and arrhythmias in TFEB overexpressing hearts. mRNA relative expression of mucolipin1 (mcoln1) gene expression in WT and TFEB/tTA mouse hearts (n=8/group) (8A), mRNA relative expression of modulatory calcineurin-interaction protein (mcip 1.4) gene expression in WT and TFEB/tTA mouse hearts (n=7/group) (8B). Western blot for mitochondrial calcium uniporter (MCU), MCU quantification by densitometry (n=6/group) (8C). Western blot for Cav1.2, Cav1.2 western blot quantification by densitometry (n=6/group) (8D-E). Representative electrocardiogram (EKG) analysis of the cardiac conduction system in WT and TEB/tTA mice 4 weeks (n=3/group) and 25 weeks (n=5/group) post doxycycline removal showing normal sinus rhythm (NSR), premature atrial contractions (PACs), atrial flutter and atrioventricular block (AV Block). *p<0.05 WT versus TFEB/tTA using Student’s T-test, graphs represent mean ± SEM.

These changes are coupled to a reduction in genes responsible for SR Ca^2+^ reuptake (atp2a2 and pln) (Fig. 6A). Atp2a2/3 encode the SERCA Ca^2+^ - ATPases (SERCA2/3) required for SR Ca2+ reuptake, while phospholamban (pln) in its unphosphorylated state inhibits SERCA2 suggesting an impairment to ER Ca^2+^ regulation (Fig. 6A). Induction of tgfb1, known to have effects on cardiac electrophysiology due to a higher frequency of spontaneous Ca^2+^ oscillations, may also contribute to arrhythmogenic phenotype in TFEB/tTA hearts (Fig. 6B). Taken together, these data indicate that in response to TFEB overexpression, calcium regulation may be impaired which could lead to lethal arrhythmias.

## Discussion

The objective of the current study was to determine the consequences of TFEB overexpression on cardiac function through its actions in canonical pathways like autophagy and mitophagy and non-canonical pathways (mitochondrial metabolism and calcium regulation). Our study shows that TFEB overexpression in cardiomyocytes leads to cardiac hypertrophy, LV dysfunction and premature death. These findings are highly significant as prior studies suggested that forced TFEB expression in the setting of advanced proteinopathy protects the heart from misfolded protein-induced cardiac proteotoxicity. AAV-9-mediated TFEB transduction resulted in 2-fold overexpression of TFEB in the myocardium and rescued advanced CryABR120G cardiomyopathy [12]. However, in previously healthy hearts, the present study identifies activation of multiple pathways that lead to maladaptive ventricular hypertrophy. Therefore, while other studies support a role for TFEB as a therapeutic strategy in cardiac proteotoxic stress, the present study identifies differential effects of TFEB in the context of cardiac pathology, which must be considered if TFEB activation is to be considered as a therapeutic strategy. The present study provides molecular insight linking TFEB overexpression in healthy cardiomyocytes and maladaptive remodeling that is mediated in part by mTOR activation, autophagy inhibition and perturbed calcium-mediated signaling.

Cardiomyocytes exit the cell cycle after birth and are terminally differentiated and therefore, do not proliferate under normal physiological conditions. Pathological hypertrophy is characterized by an increase in cardiomyocyte size and thickening of the ventricular walls leading to systolic and diastolic dysfunction and heart failure [29]. We observed an increase in cardiac hypertrophy with forced TFEB expression compared to WT mice after 1- and 2-weeks which correlated with reduced LV function (HFrEF) after 2-weeks. These findings are supported by an increase in positive regulators of hypertrophy genes after 1 and 2-weeks. Another hallmark of persistent cardiac hypertrophy is irreversible maladaptive changes, leading to fibrosis [30]. Cardiac fibrosis involves the deposition of collagen rich extracellular matrix in the interstitial space. The absence of phenotypic evidence of fibrosis in TFEB overexpression hearts using Picrosirius Red staining (PSR) and Masson’s Trichrome staining after two weeks, and the normalization of transcriptomic signatures of fibrosis that were evident at one week but resolved by two-weeks suggest that TFEB overexpression could be associated with active remodeling of the extracellular matrix.

Autophagy induction is one of the mechanisms by which TFEB induction mediates therapeutic benefit in certain pathophysiological states. During cardiac ischemia, an increase in autophagy can be protective [31]. Conversely, autophagy activation can be detrimental in pressure overload-induced heart failure [32], or in a model of ischemia/reperfusion [32]. Other studies support a protective role of increased autophagy in ATG5 heart KO mice [33] and in humans with dilated cardiomyopathy [34]. Inducing autophagy in a model of cardiac proteotoxicity (CryAB^R120G^ mutation) by overexpressing ATG7, increased autophagic activity [35] and protected the heart from protein aggregation. Using the same model, Bo Pan et al. [11] demonstrated that viral mediated TFEB overexpression in neonatal rat ventricular myocytes (NRVMs) reduced CryAB^R120G^ positive aggregates by inducing autophagy. Using an autophagic flux assay and measuring the relative turnover of LC3-II in the presence and absence of the lysosomal inhibitor, chloroquine (CQ), we observed no further accumulation of LC3-II or any change in the LC3-II/I ratio, which was contrary to previous reports and might have been the result of the supra-physiological TFEB levels. mTOR is recruited to the lysosomal membrane by Rag GTPases and is a known inhibitor of autophagy. A recent study found that MiT/TFE transcription factors, including TFEB, control the mTOR lysosomal recruitment and activity by directly regulating the expression of Rag D in kidney cells [36]. The silencing of Rag D reversed mTOR hyperactivation. Thus, our studies support a model in which in the healthy heart overexpression of TFEB inhibits autophagic flux, potentially in part by persistent activation of mTOR. In our model, inhibition of mTOR after 4-weeks of rapamycin treatment partially improved cardiac function in TFEB hearts relative to mice treated with vehicle, emphasizing the possibility of a negative feedback loop whereby TFEB upregulates mTOR. However, our data indicate that mTOR is only partially responsible for the contractile dysfunction and pathological hypertrophy in TFEB/tTA mice. Moreover, selective autophagy, which targets damaged mitochondria to the lysosome for degradation is termed, mitophagy, and was impaired in TFEB/tTA hearts compared to WT. Several studies have reported that impaired mitophagy exacerbates stress induced heart failure [37–39] while PINK1 levels are known to be repressed in humans with heart failure [38]. Thus, autophagy impairment might contribute to the maladaptive phenotypes with TFEB is activated in the normal heart, by impairing mitochondrial quality control.

Mitochondria are critical for energy generation and are involved in reactive oxygen species generation, signal transduction, Ca^2+^ regulation and cell death [40]. Mitochondrial dysfunction is a central characteristic of heart failure. Balanced fission and fusion (mitochondrial dynamics) are important to preserve mitochondrial integrity and cardiac function. Mitochondrial fusion is regulated by the GTPases OPA1, MFN1 and MFN2 whereas DRP1 and Fis1 mediates fission. TFEB overexpression in skeletal muscle by adeno associated virus (AAV-TFEB) resulted in increased mitochondrial number, size and mitochondrial DNA, which supports a role for TFEB in mitochondrial biogenesis in skeletal muscle [13]. We observe altered mitochondrial morphology with an increase in mitochondrial number and reduced cristae index score. Despite an increase in the direct TFEB target, PGC1alpha protein and gene expression after 2-weeks, the net effect on mitochondrial proteins at 1-week is repression. Moreover, PGC-1α upregulates and activates TFEB [41] suggesting the presence of pro-survival feedback loop [8]. Fatty acid is the preferred substrate in the adult myocardium and supplies 70% of total ATP in a normal heathy heart. FA utilization is reduced in HF [42], while cardiac glucose utilization is increased initially and ultimately decreased as systolic dysfunction progresses [43] in parallel with transcriptional repression of genes that regulate these processes. The observed reduction in gene expression in FA genes, oxidative phosphorylation genes and pyruvate metabolism genes reveal an important effect of TFEB in mediating the transcriptional repression of mitochondrial metabolic pathways in the heart. After 2-weeks, mitochondrial protein expression was also reduced. Despite this, mitochondrial respiration and ATP production remains unchanged. These findings contrast with Mansueto et al. (2017), who demonstrated that TFEB overexpression in skeletal muscle induced mitochondrial biogenesis, improved mitochondrial respiratory complex activity and ATP production in a PGC-1α independent manner. Given the established role of TFEB in mediating lysosomal biogenesis and considering that autophagy is inhibited in our model, the possibility is raised that an imbalance between lysosomal activation combined with autophagy inhibition to mimic characteristics of lysosomal storage disorders [44].

Recent discoveries of interconnections between the mitochondria and the endoplasmic reticulum (ER) membrane provide novel perspectives for understanding molecular events in cardiac pathophysiology. Moreover, the ER is a central site for Ca^2+^ storage and homeostasis [45]. Alterations in luminal Ca^2+^ can lead to protein misfolding and subsequent activation of the unfolded protein response (UPR). It is now well established that altered mitochondrial function and dynamics can lead to ER stress [46]. In the context of cardiac hypertrophy and reduced mitochondrial gene and protein expression in TFEB overexpression hearts, we also observe an increase in the UPR as evidenced by an increase in IRE1α, PERK and p-eif2α. Altered Ca^2+^ homeostasis may exacerbate mitochondrial alterations. Additionally, calcium dysregulation could contribute to development of lethal arrhythmias in TFEB overexpressing mice [47]. Moreover, deranged Ca^2+^ homeostasis could contribute to the progression of LV dysfunction [48]. Altered calcium dynamics could also be responsible for increased lysosomal calcium release and activation of calcineurin in TFEB/tTA mice which could contribute to pathological cardiac hypertrophy and heart failure [49].

In summary, the present study identifies multiple mechanisms by which TFEB overexpression in the healthy heart induces adverse LV remodeling and contractile dysfunction. Proximal mechanisms include broad suppression of mitochondrial energy metabolic pathways and perturbed regulation of calcium signaling. Although reversible in the short term, these data indicate that for TFEB modulation to be adopted as a therapeutic strategy for cardiac pathology, it might need to be activated in an intermittent manner to mitigate potential adverse consequences.

## Abbreviations

AAV: Adeno-associated viruses
α-MHC: alpha- myosin heavy chain
CMV: Cytomegalovirus
CQ: Chloroquine
DMSO: dimethyl sulfoxide
DTNB 5: 5-dithio-bis-(2-nitrobenzoic acid
E-C: coupling Excitation-Contraction coupling
EDV: End-Diastolic Volume
EF: Ejection Fraction
EKG: electrocardiogram
ER: Endoplasmic Reticulum
ESV: End-Systolic Volume
HFrRF: Heart Failure reduced Ejection Fraction
LC3B: Microtubule-associated protein light chain
LV: Left ventricle
MAPK1: Mitogen-activated protein kinase 1
MCOLN1: Mucolipin 1
MCU: mitochondrial calcium uniporter
MFN: Mitofusin
Mit/TFE: microphthalmia/transcription factor E
NADP^+^: nicotinamide adenine dinucleotide phosphate
OPA1: Optic atrophy 1
PFA: paraformaldehyde
PGC1α: Peroxisome proliferator activated receptor-γ coactivator 1α
PVDF: Polyvinylidene fluoride or polyvinylidene difluoride
RIPA: Radioimmunoprecipitation assay buffer
SR: Sarcoplasmic reticulum
STED: Stimulated emission depletion
TAC: Transaortic constriction
TetO: Tetracycline Operator
TFEB: Transcription factor EB
TRE: Tetracycline response element
tTA: Tetracycline
tTA: tetracycline-controlled transactivator protein
UPR: unfolded protein response
WGA: Wheat Germ Agglutinin

## Acknowledgements

The authors thank Chantal Allamargot (University of Iowa Microscopy Core, University of Iowa) and Kathy Zimmerman (Imaging core, University of Iowa) and Jamie Soto (Metabolic Phenotyping Core, University of Iowa) for their technical assistance. The Mito-Keima mice were a kind gift from Dr. Torn Finkel (Center for Molecular Medicine, NHLBI, NIH). RNA-seq data were obtained at the Genomics Division of the Iowa Institute of Human Genetics, which is supported, in part, by the University of Iowa Carver College of Medicine. This work was supported by NIH grants 1R01 HL112413, R01HL108379 to EDA and the AHA award 16SFRN31810000 to EDA who is an established investigator of the AHA.

## Author Information

**Division of Endocrinology and Metabolism, Department of Internal Medicine, University of Iowa Hospitals and Clinics, University of Iowa, Iowa City, IA.**

Helena C. Kenny, Eric T. Weatherford, Greg C. Collins, Taha Gesalla, Harsh Goel, Theo Romac, Renata Pereira, Jennifer Streeter, Jacob Sharafuddin, Long-Sheng Song, E. Dale Abel

**Fraternal Order of Eagles Diabetes Research Center, University of Iowa.**

Helena C. Kenny, Eric T. Weatherford, Greg C. Collins, Taha Gesalla, Harsh Goel, Theo Romac, Renata Pereira, Jennifer Streeter, Jacob Sharafuddin, E. Dale Abel

**Abboud Cardiovascular Research Center, Carver College of Medicine, University of Iowa, Iowa City, IA, USA**

Kathy Zimmerman, Jared M. McLendon, Long-Sheng Song

**Central Microscopy Research Facility, University of Iowa**

Chantal Allamargot

**Department of Pathology, University of Iowa**

Dao-Fu Dai

**Center for Cardiovascular Research and Division of Cardiology, Washington University School of Medicine.**

Dr. Abhinav Diwan

Ethics Declaration

## Author Contributions

HK and EDA designed the research study. ETW, GC, CA, TG, KZ, HG, JMM, TR, JS and DFD performed the research. JS reviewed all electrocardiograms, AD and LSS provided materials and methodology support. HK, CA and ETW analyzed the data. RP and LSS provided feedback on the writing. HK and EDA wrote the manuscript.

