## Supplemental Material for "Cardiac Specific Overexpression of Transcription Factor EB (TFEB) in Normal Hearts Induces Pathologic Cardiac Hypertrophy and Lethal Cardiomyopathy"

**Supp. Table 1.**

| Antibody Name | Company | Product Number | Molecular Weight (KDa) |
|---|---|---|---|
| TFEB | Bethyl | A303-673A | 65 |
| TFEB | Santa Cruz | SC 48784 | 65 |
| p-mTOR S448 | Cell signaling | CS 2971S | 289 |
| mTOR | Cell signaling | CS L27D4 | 289 |
| p-S6 S235/236 | Cell signaling | CS 2211S | 32 |
| S6 | Cell signaling | CS 2317S | 32 |
| GAPDH | Cell signaling | CS 2118L | 37 |
| p-ULK1 Ser757 | Cell signaling | CS 6888S | 140 |
| ULK1 | Cell signaling | CS 8054 | 140 |
| LC3B | Cell signaling | CS 41085 | 14 &16 |
| P62/DNM1L | Cell signaling | CS 5114S | 62 |
| Mfn1 | Abcam | ab 57602 | 75 |
| Mfn2 | Abcam | Ab50843 | 75 |
| OPA1 | Biological sciences | BD 612606 | 80-100 |
| YME1L | Protein Tech | 11510-1-AP | 60 |
| OMA1 | Sigma | SAB2108766 | 37 |
| DRP1/DNM1L | Abnova | H00010059 | 50 & 70 |
| NDUFa9 (CI) | Abcam | ab 14713 | 36 |
| SDHa (CII) | Abcam | ab 14715 | 70 |
| MTCO3 (CIII) | Abcam | ab 14705 | 40 |
| UQCRC1 (CIV) | Abcam | ab 110252 | 53 |
| Atp5a (CV) | Abcam | ab 14748 | 55 |
| MCU | Sigma | HPA 016480 | 27 |
| P-eif2alpha | Cell Signaling | CS 35975 | 38 |
| Eif2alpha | Santa Cruz | 81201 | 38 |
| IRE1alpha | Cell Signaling | 32945 | 130 |
| GRP78/Bip | BD Bioscience | BDB 610978 | 75 |
| Cathespin B | Cell Signaling | 31718S | 44 |
| Cav1.2 | Abcam | Ab58552 | 249 |

**Supp. Table 2.**

| Primer Name | Forward Sequence | Reverse Sequence |
|---|---|---|
| Tfeb | AAG GAG CGG CAG AAG AAA GA | CCT TGA GGA TGG TGC CTT TG |
| Nppa | ATG GGC TCC TTC TCC ATC A | CCT GCT TCC TCA GTC TGC TC |
| Nppb | GGA TCT CCT GAA GGT GCT GT | TTC TTT TGT GAG GCC TTG GT |
| Acta1 | CCTGTATGCCAACAACGTCA | CTCGTCGTACTCCTGCTTGG |
| Myh6 | AGA TAG TGG AAC GCA GGG ATG | CTT CAG CAG CGG TTT GAT CTT G |
| Myh7 | CTCCTGCTGTTTCCTTACTTGC | CTCCTGCTGTTTCCTTACTTGC |
| Col11a1 | ACA GTA GCA CAA ACA GAG GCA A | AAT CCC TGC CGT CTA CTC CT |
| Col11a2 | ATC AGT ACG AAA GGG CCC CA | TAA ACC CAT TGG TCC AGG GC |
| Col14a1 | TGA AGC ACC CAC AGC CAT AG | TCC AGG CAC CAT AAC CGT TC |
| Col1a1 | AGA TGT AGG AGT CGA GGG AC | GGC CTT GGA AAC CTT GTG GA |
| Col5a | TTT GGA CGA GTA GGG CAA CC | GTT TCC AGG GGG ACC CTT TT |
| Col3a1 | GTC CAA CTG GTG GCC AAA ATT A | TTG CGT CCA TCA AAG CCT CT |
| Tgfbr1 | CCT TCA AAC GCG CTG ACA TC | CCA TCA CTC TCA AGG CCT CA |
| Tgfb2 | TCC CCT CCG AAA ATG CCA TC | AGG TGC CAT CAA TAC CTG CAA |
| Atp7a | GGA AAC CTA CTT TCC CGG CT | GGG TTC CAT GGG TGA TGG TT |
| GAPDH | AAC GAC CCC TTC ATT GAC | TCC ACG ACA TAC TCA GCA C |

### Supplemental figure 1. Pathological cardiac hypertrophy develops following TFEB induction in cardiomyocytes.

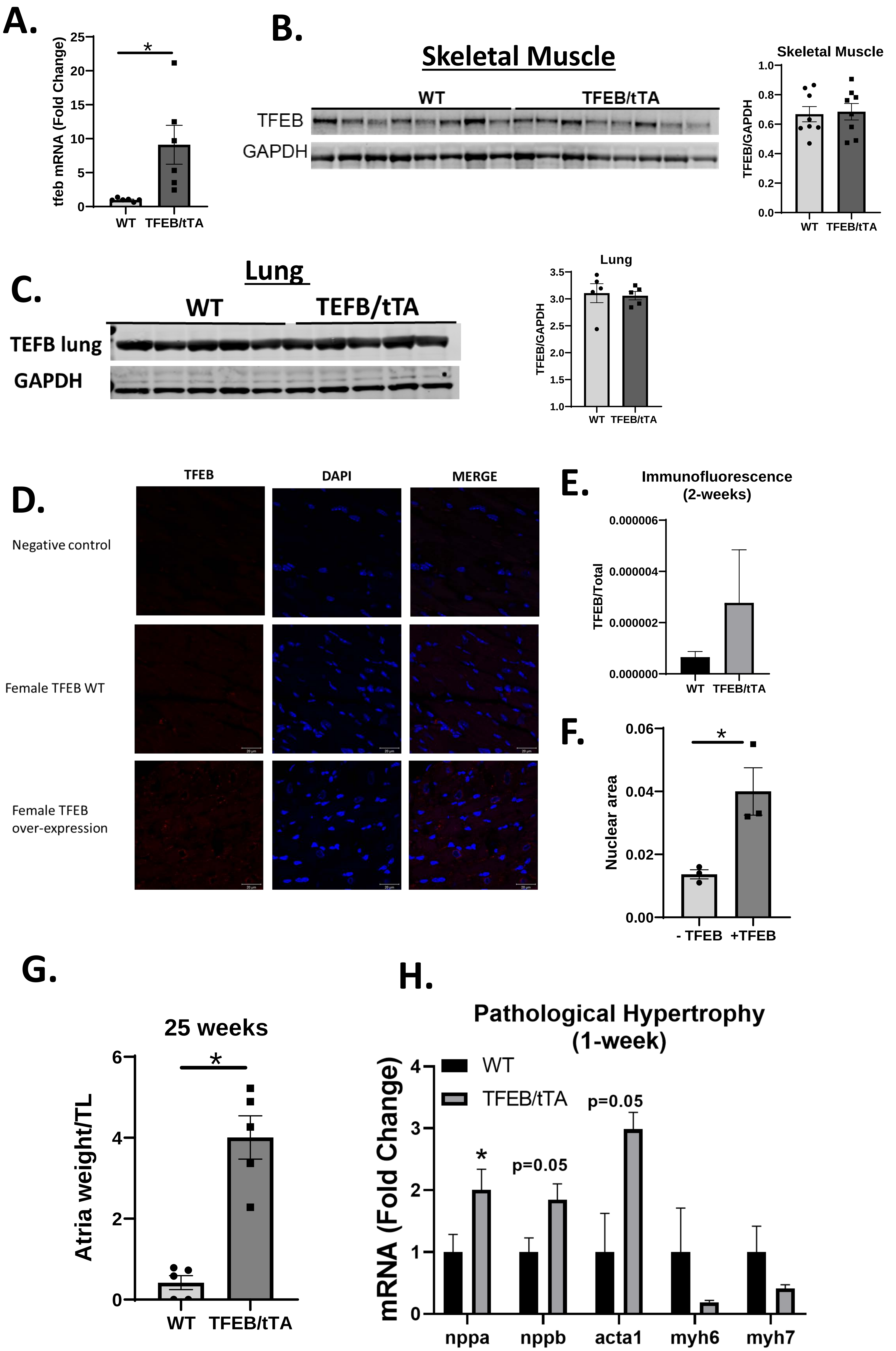

**Supplemental Figure 1 Pathological cardiac hypertrophy develops following TFEB induction in cardiomyocytes.**

A: mRNA fold change of tfeb in the heart 2-weeks following doxycycline removal (n=6/group)

B-C: TFEB immunoblot in skeletal muscle and lung tissue in WT and TFEB/tTA mice, quantification by densitometry (2-weeks) following doxycycline withdrawal

D-F: Immunofluorescence images for TFEB (red), nuclei (dapi, blue) and merged images, in WT and TFEB/tTA (2-weeks) after doxycycline removal, negative control; same as experimental samples except no TFEB antobdy, TFEB immunofluorescence quantification of pixel intensity (TFEB/total) and nuclear area of nuclei with and without TFEB accumulation.

G: Atrial weight/tibia length after 25 weeks of TFEB induction (n=5/group).

H: mRNA fold change for markers of pathological hypertrophy after 1-week TFEB induction (nppa, nppb, acta1, myh6, myh7).

Graphs represent mean ± SEM. P<0.05 compared to WT

### Supplemental figure 2. Heart failure and premature mortality following TFEB overexpression in cardiomyocytes

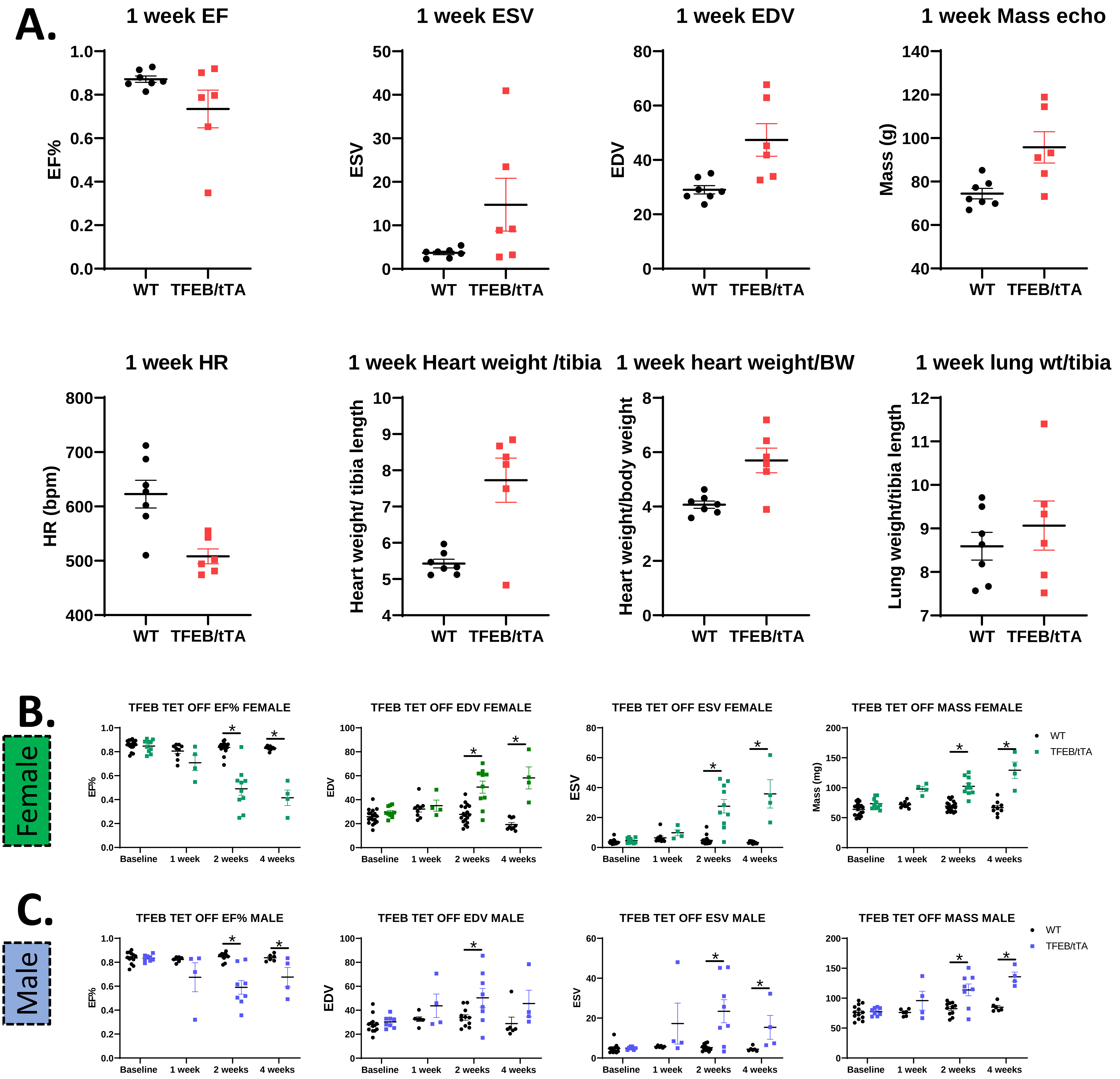

**Supplemental Figure 2. Heart failure and premature mortality following TFEB overexpression in cardiomyocytes**

A: echocardiography data depicting ejection fraction (EF%), end-systolic volume (ESV), end-diastolic volume (EDV), mass, heart rate (HR) in beats per minute (bpm) after 1-week and heart weight/tibia length, heart weight/body weight and lung weight/tibia length (1-week) after doxycycline withdrawal.

B-C: Echocardiographic analysis in male and female mice; ejection fraction (EF%), end-diastolic volume (EDV), end-systolic volume (ESV), mass, heart rate (HR) in beats per minute (bpm) after 2-weeks after doxycycline withdrawal.

Graphs represent mean ± SEM. P<0.05 compared to WT

### Supplemental figure 3. TFEB overexpression and cardiac fibrosis after 1- and 2-weeks doxycycline removal

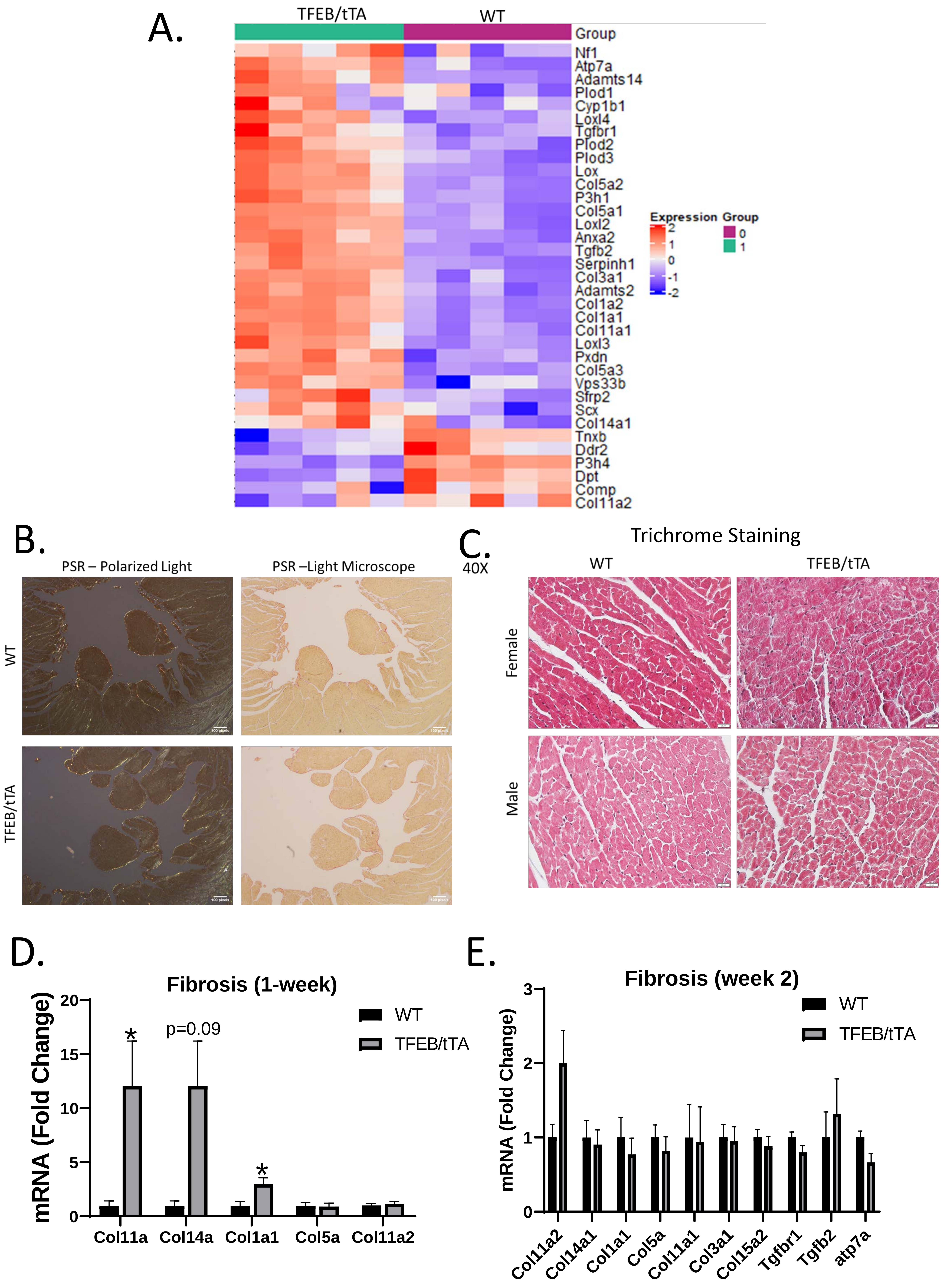

**Supplemental Figure 3. TFEB overexpression and cardiac fibrosis after 1- and 2-weeks doxycycline removal**

A: Heatmap of log2(x+1) normalized counts of differentially expressed genes involved in fibrosis identified by enrichment analysis performed using Enrichment Browser and the Pathway Analysis with Down-weighting of Overlapping Genes (PADOG) algorithm.

B: Representative images of Picosirus Red staining using polarized and light microscope in WT and TFEB/tTA paraffin embedded heart sections at 2-weeks, scale bar 100 pixels

C: Representative images of trichrome staining in WT and TFEB/tTA paraffin embedded heart sections at 2-weeks, scale bar 20µM

D-E: qPCR analysis for markers of fibrosis in 1- and 2-week samples

Graphs represent mean ± SEM. P<0.05 compared to WT

### Supplemental figure 4. Reversibility of heart failure following reversal of TFEB overexpression

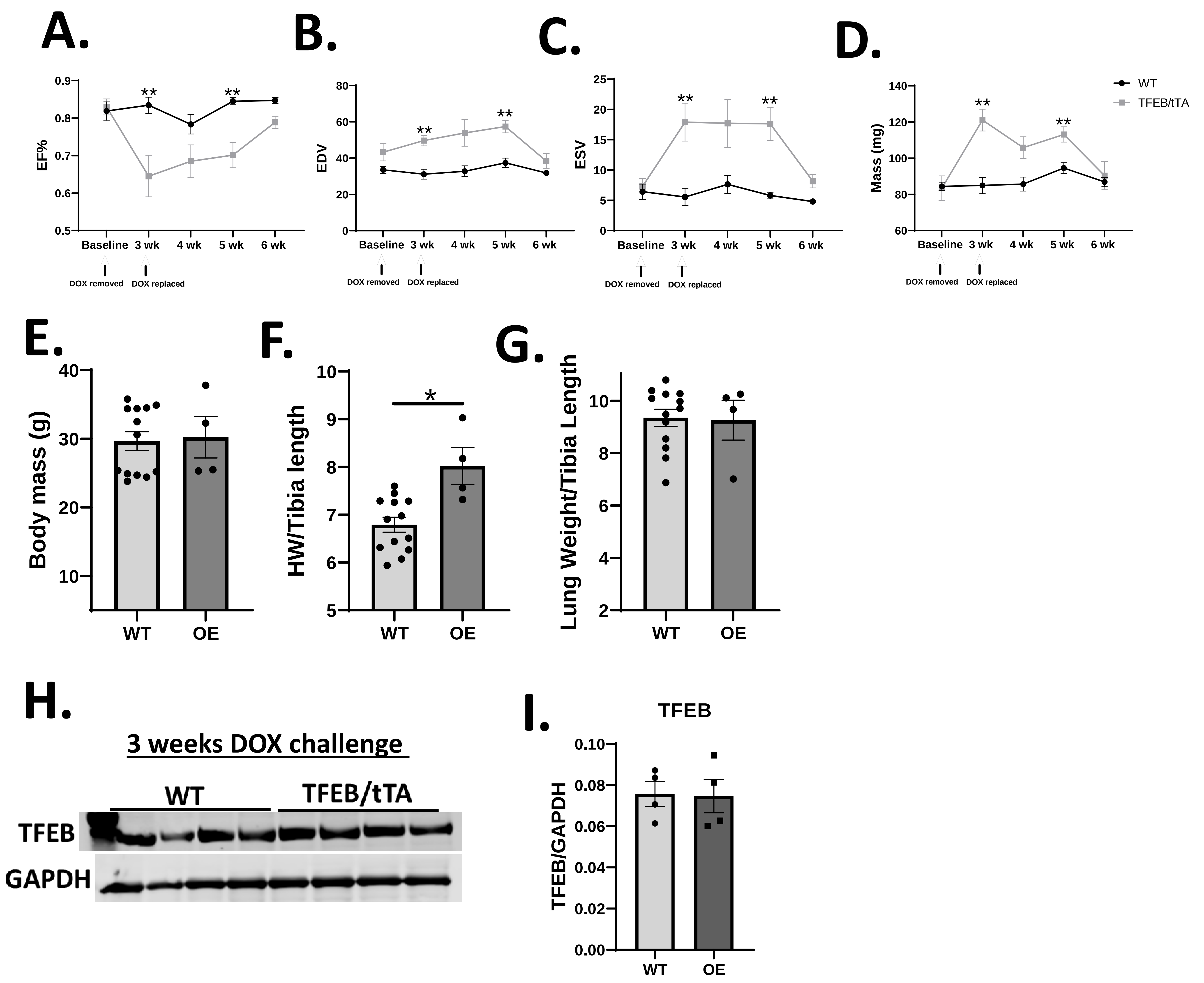

**Supplemental Figure 4. Reversibility of heart failure following reversal of TFEB overexpression**

A-D: Echocardiographic analysis on WT and TFEB/tTA mice at baseline, 3-weeks after doxycycline removal, at 4-weeks, 5-weeks and 10-weeks, ejection fraction (EF%), end-diastolic volume (EDV), end-systolic volume (ESV), mass.

E-G: body mass, heart weight/tibia length and lung weight/tibia length at tissue harvest

H-I: TFEB immunoblot image and quantification in WT and TFEB/tTA after 3 weeks of TFEB induction (-DOX) and 3-weeks TFEB suppression (+ DOX). Tissue harvested at 6 weeks

Graphs represent mean ± SEM. P<0.05 compared to WT

### Supplemental figure 5. TFEB overexpression in cardiomyocytes inhibits autophagy and mitophagy.

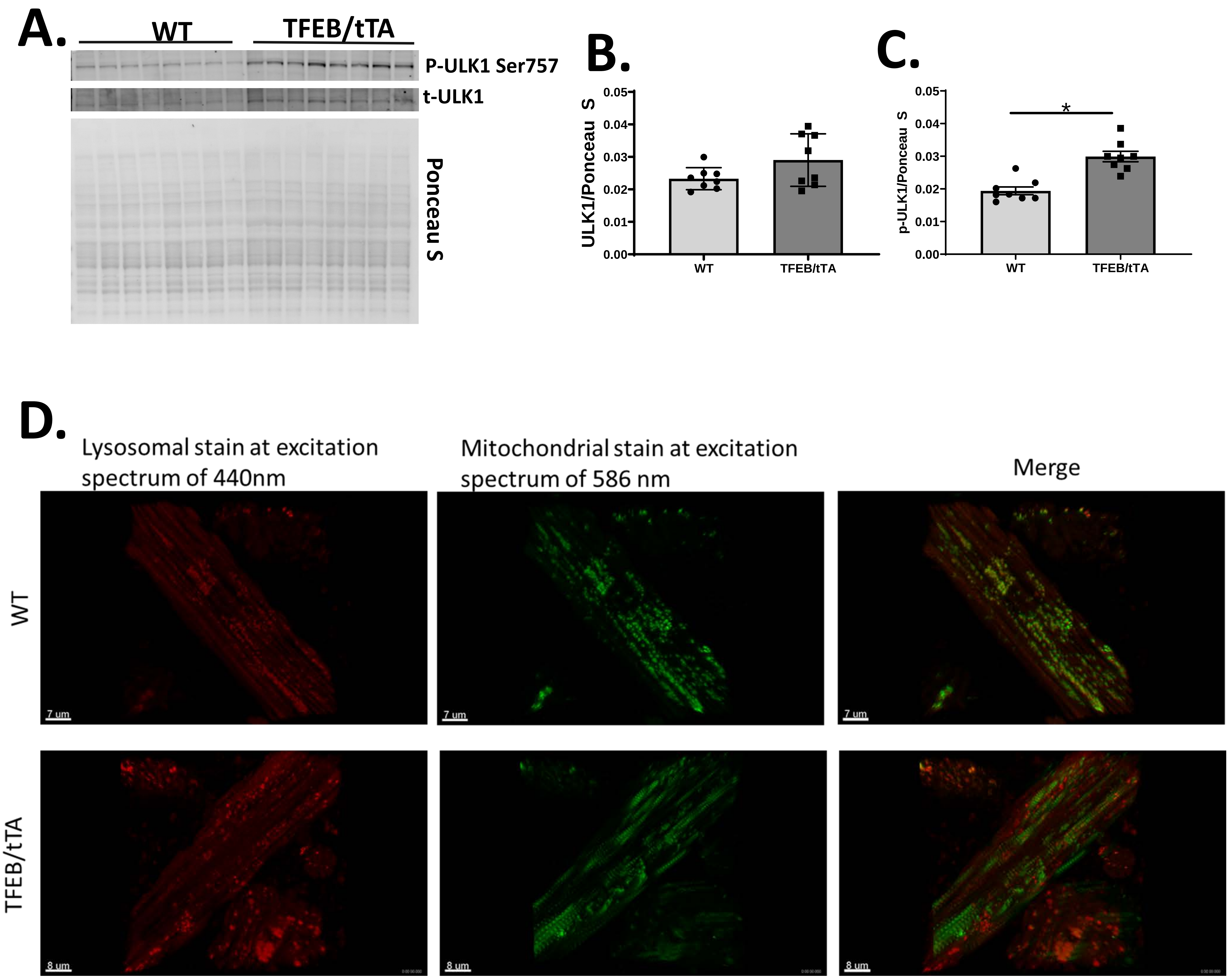

**Supplemental Figure 5 TFEB overexpression in cardiomyocytes inhibits autophagy and mitophagy.**

A-C: Western blot images showing phosphorylated ULK1 at Ser757 (p-ULK1) and total ULK1 (t-ULK1) (t-ULK1) in WT and TFEB/tTA heart lysates after 2-weeks (WT n=8, TFEB/tTA n=8). Quantification of western blots by densitometry and normalized to total protein (ponceau S).

D: WT and TFEB/tTA mice expressing the fluorescent protein keima at an excitation spectrum (458nm) (lysosomal) and longer wavelength (561nm).

Student's T-test and two-way ANOVA were used for statistical analysis, graphs represent mean ± SEM.

Graphs represent mean ± SEM. P<0.05 compared to WT

Supplemental figure 6. mTOR signaling is induced in TFEB overexpression hearts after 1-week

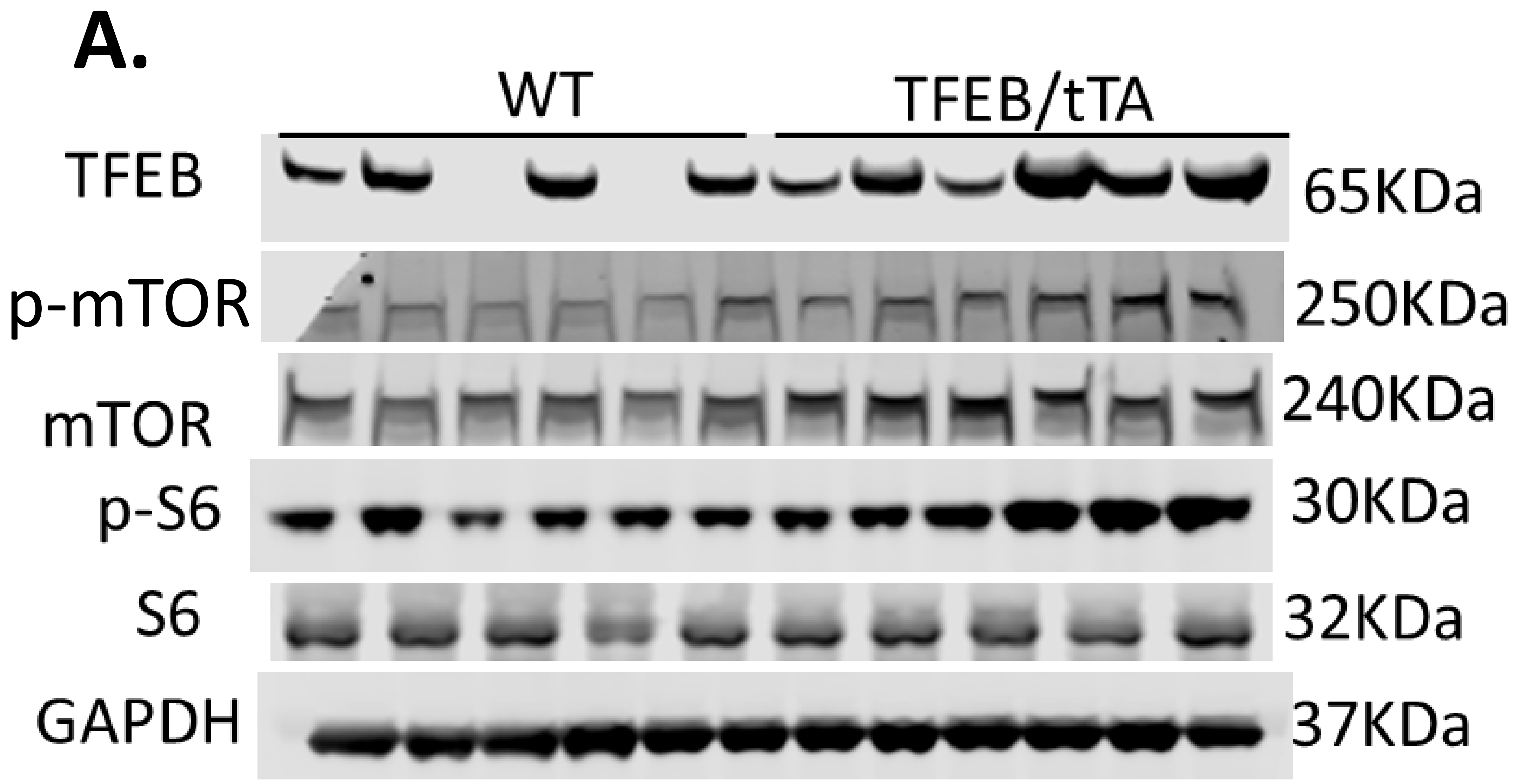

**Supplemental Figure 6. mTOR signaling is induced in TFEB overexpression hearts after 1-week**

A: Immunoblot images after 1-week in WT and TFEB/tTA heart tissue (n=6/group) for TFEB, p-mTOR, mTOR, p-S6, S6 and GAPDH

### Supplemental figure 7. RNA-seq analysis of mRNA isolated from mice following overexpression of TFEB for one week

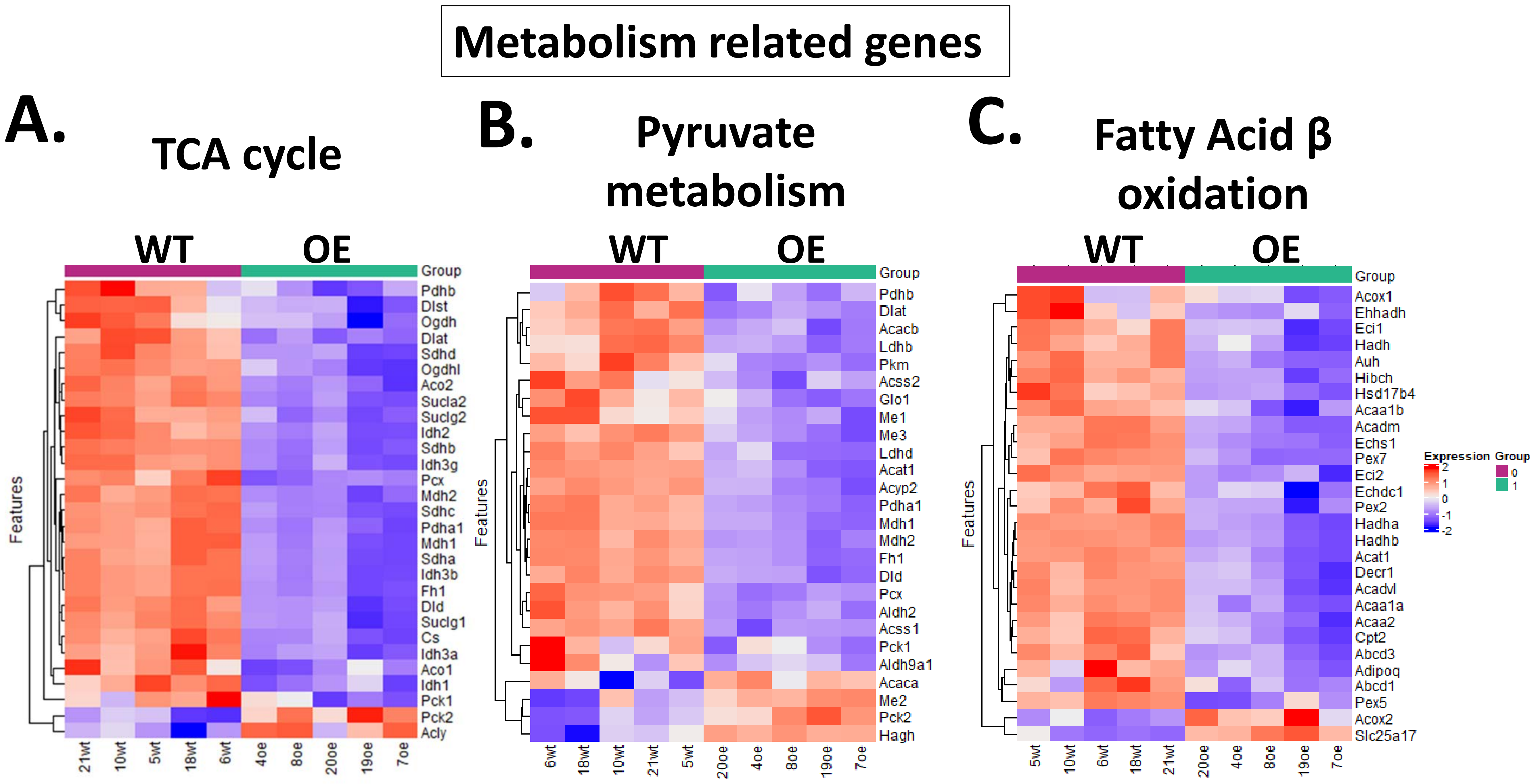

**Supplemental Figure 7. RNA-seq analysis of mRNA isolated from mice following overexpression of TFEB for one week**

A-C: Heatmap of log2(x+1) normalized counts of differentially expressed genes involved in fibrosis identified by enrichment analysis performed using Enrichment Browser and the Pathway Analysis with Down-weighting of Overlapping Genes (PADOG) algorithm depicting gene changes for tricarboxylic acid cycle (TCA) (A), pyruvate metabolism (B) and fatty acid β oxidation (C). Gene changes were filtered based on p value of $<0.05$.

### Supplemental figure 8. Mitochondrial morphology and function following 2 weeks of TFEB overexpression in the heart.

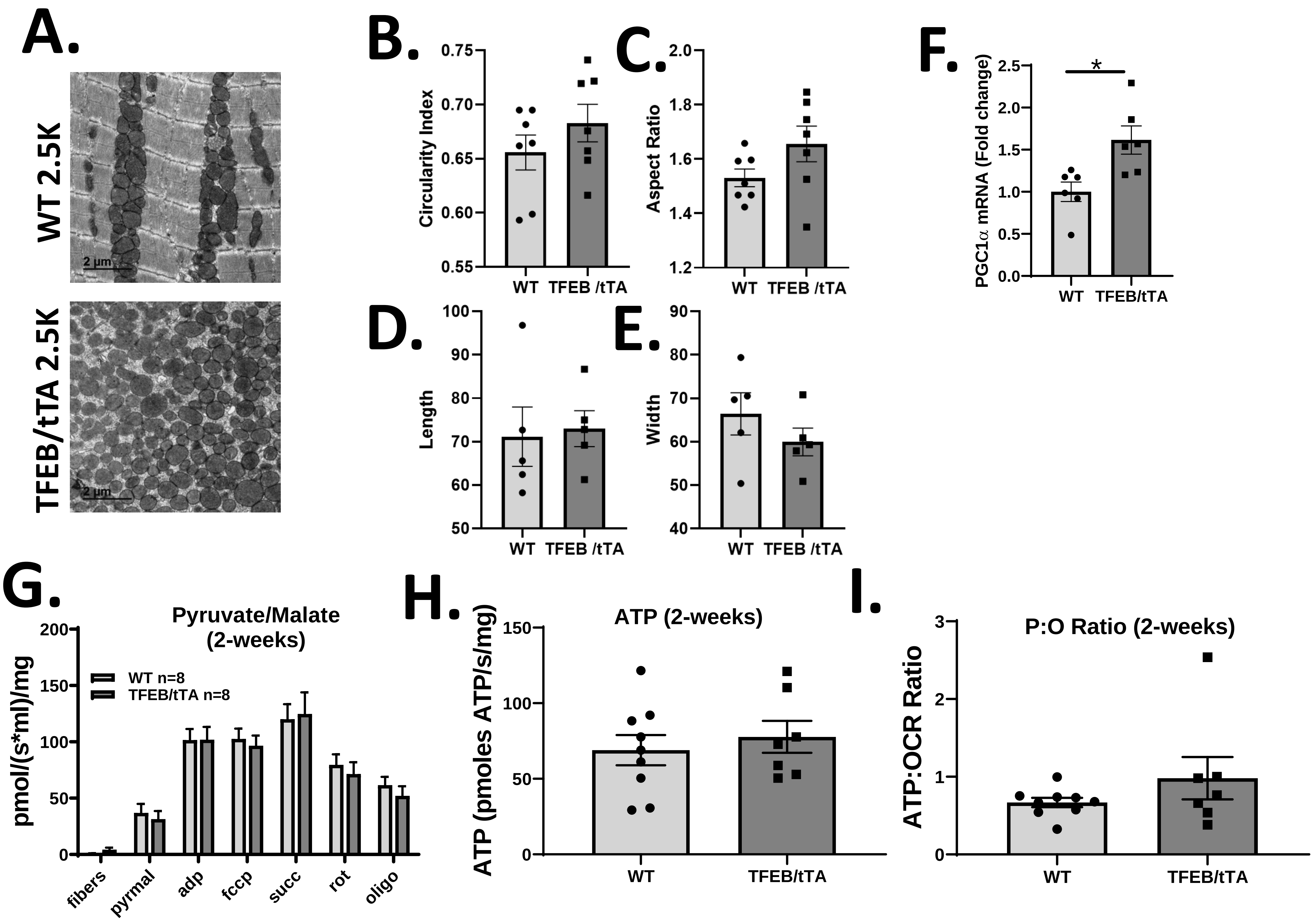

**Supplemental Figure 8 Mitochondrial morphology and function following 2 weeks of TFEB overexpression in the heart.**

A-E: Tem images in WT and TFEB/tTA hearts (magnification 2500X and scale bar 2µm), quantification of TEM; circularity index, aspect ratio, length and width.

F: PGC1α mRNA fold change in WT and TFEB/tTA hearts (n=6/group).

G-I: Mitochondrial respiration with pyruvate/malate as substrates in WT and TFEB/tTA permeabilized heart fibers (n=8/group), ATP production in WT and TFEB/tTA permeabilized heart fibers (n=8/group), P:O ratio WT and TFEB/tTA permeabilized heart fibers (n=8/group)

Graphs represent mean ± SEM. $P<0.05$ compared to WT

### Supplemental figure 9. TEFB overexpression results in ER stress signaling after 2-weeks doxycycline withdrawal

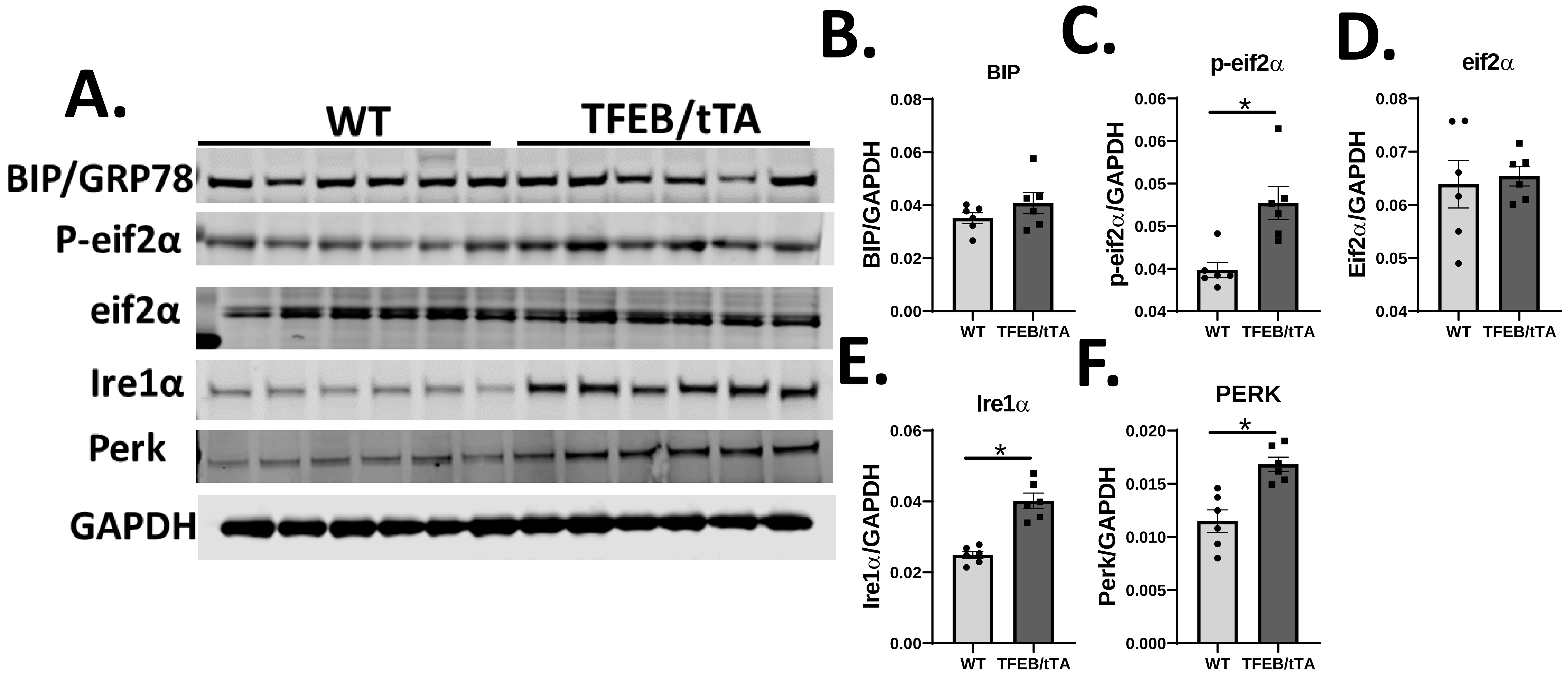

**Supplemental Figure 9. TEFB overexpression results in ER stress signaling after 2-weeks doxycycline withdrawal**

A-F: Western blot images for markers of endoplasmic reticulum (ER) stress (TFEB, p-eif2alpha, Grp78, IRE1alpha, GAPDH) in WT and TFEB/tTA heart lysates (n=6/group), quantification by densitometry.

Graphs represent mean ± SEM. P<0.05 compared to WT

Supplemental figure 10. Cardiac arrhythmias in TFEB overexpressing hearts.

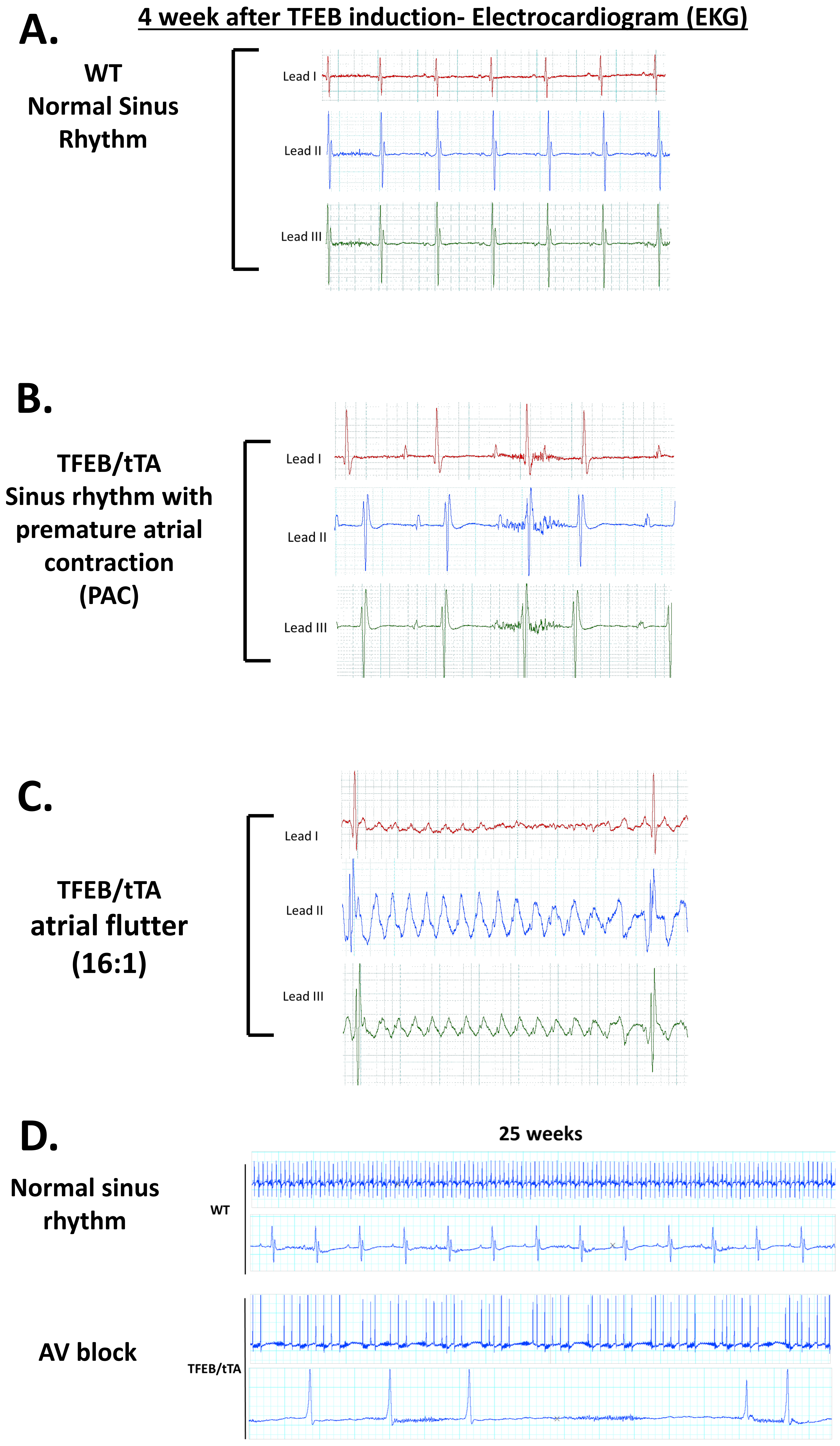

**Supplemental Figure 10. Cardiac arrhythmias in TFEB overexpressing hearts.**

A-D: Representative electrocardiography (EKG) traces from lead I, II and III for WT demonstrating normal sinus rhythm, premature atrial contractions (PACs) and atrial flutter, 4-weeks after TFEB induction in  and TFEB/tTA mice (PACs and atrial flutter) after 4 weeks TFEB induction and atrioventricular block observed 25 weeks TFEB induction.
